# A new stress-response pathway in *Mycobacterium tuberculosis*

**DOI:** 10.64898/2026.04.15.718621

**Authors:** Pieter M.M. van der Velden, Jona Merx, Lars van Dijk, Femke Roos, Julia Brake, Tom Berben, Paul B. White, Rob M. de Graaf, Eva Terschlüsen, Ingrid J.M. van Weerdenburg, Bai H.E. Zhang, Reinout van Crevel, Laura van Niftrik, Jakko van Ingen, Thomas J. Boltje, Robert S. Jansen

## Abstract

Metabolic adaptations are key in the virulence of the pathogen *Mycobacterium tuberculosis* (Mtb). However, our current understanding of these adaptations is limited to common pathways in central carbon metabolism. Here we apply an untargeted bottom-up approach to discover unknown stress-responsive metabolites and their biosynthetic enzyme. We show that upon exposure to hypoxia, nitric oxide and activated macrophages, Mtb rapidly produces high levels of an unknown metabolite that we identify as 6’-γ-amino-butyric acid-trehalose (GABA-trehalose). Formation of millimolar GABA-trehalose levels under these stresses is driven by a rapid rise in GABA, which is also excreted. We demonstrate that GABA-trehalose is produced from GABA and trehalose by the uncharacterized ATP-grasp enzyme Rv1722, involving a carboxylate-hydroxyl ligation that is non-canonical for ATP-grasp enzymes. Phylogenetic analyses demonstrate that the gene *rv1722* is present in most slow-growing mycobacteria but absent in most rapid-growing mycobacteria. While the role of GABA-trehalose in Mtb metabolism remains unclear, we postulate that the increased NADH/NAD+ ratio under hypoxia and nitric oxide exposure promotes GABA formation and inhibits its breakdown, leading to GABA accumulation and excretion. Rv1722-driven coupling of GABA and trehalose constitutes an alternative to excretion that conserves carbon and nitrogen. Taken together, our bottom-up approach reveals a new stress-response pathway in Mtb that rapidly produces large quantities of GABA-trehalose. These findings extend our knowledge of the metabolic adaptations that a major human pathogen utilizes in response to immune system-imposed stresses.

## Introduction

Tuberculosis has plagued humans for thousands of years and still kills over a million people annually(1). Over these years, *Mycobacterium tuberculosis* (Mtb) has evolved into a professional pathogen that uses humans as its sole reservoir and defies our immune system. Metabolic adaptations have been shown to be key in this evolution by enabling the use of host-derived nutrients and thwarting chemical attacks launched by our immune system(2–4).

One of harshest environments encountered by the intracellular pathogen Mtb is the phagolysosome of human macrophages. Here, Mtb is exposed to a combination of hypoxia, reactive nitrogen species, reactive oxygen species and low pH(5). Metabolomics has been key in the discovery of distinct metabolic strategies that Mtb employs to survive these stresses(6–8).

While technical advances in metabolomics have rapidly increased our understanding of Mtb metabolism, current knowledge is still largely based on metabolites involved in classical biochemical pathways, such as glycolysis and the tricarboxylic acid (TCA) cycle(9). At the same time, untargeted metabolomics studies have revealed that large parts of metabolomes are uncharacterized and that even in the best characterized organisms, many metabolites remain unannotated(10–12). Therefore, we hypothesized that metabolic adaptation in Mtb likely involves previously unidentified metabolites. Though challenging, the discovery of new metabolites has often led to entirely new fields of investigation(13–16).

To extend our knowledge on metabolic adaptations in Mtb from classical biochemical pathways into unknown chemical space, we applied a bottom-up approach in which we first screened for unknown stress-responsive metabolites and then characterized their biosynthetic enzyme and related metabolic pathway. Here, we identify GABA-trehalose as a new, highly abundant, stress-responsive metabolite in Mtb. This novel metabolite is rapidly formed from GABA and trehalose by the atypical ATP-grasp enzyme Rv1722.

## Materials & Methods

### Mycobacterial strains & culturing

*M. tuberculosis* mc^2^ 6206 H37Rv::ΔpanCD, ΔleuCD was kindly provided by Dr William Jacobs (Albert Einstein College of Medicine, New York, USA). The strains *M. smegmatis* mc^2^ 155, *M. abscessus* ATCC 19977, *M. kansasii* ATCC 12478, and *M. bovis* BCG Danish were provided by the mycobacteriology department of the Radboud University medical center (Nijmegen, the Netherlands). All strains except wildtype *M. tuberculosis* H37Rv were cultivated at 37°C in Middlebrook 7H9 supplemented with albumin, dextrose and sodium chloride (7H9-ADN) or on Middlebrook 7H10-ADN. 7H9-ADN liquid medium was prepared with 0.47% w/v dehydrated 7H9 broth (Becton, Dickinson and Company (BD), Franklin Lakes, USA), 0.2% v/v glycerol, 0.5% w/v fatty acid-free bovine serum albumin, 0.2% w/v D-glucose, 0.085% w/v NaCl, and 0.04% v/v Tyloxapol. 7H10-ADN solid medium was prepared with 1.9% w/v dehydrated Middlebrook 7H10 broth (BD), 0.5% w/v fatty acid-free bovine serum albumin, 0.2% w/v D-glucose, and 0.085% w/v NaCl. Solid and liquid media for *Mtb* mc^2^ 6206 H37Rv::ΔpanCD, ΔleuCD were supplemented with 24 μg/mL calcium pantothenate and 50 μg/mL L-leucine. Wildtype *M. tuberculosis* H37Rv was grown in liquid 7H9 media with oleic acid-albumin-dextrose-catalase (OADC) supplement. Small volume cultures (≤40 mL) were not shaken, while larger volume cultures were shaken at 90 rpm.

### Mycobacterial stress exposure

Biomass equivalent to 1 mL of culture at an optical density at 600nm (OD_600_) of 1.0 was vacuum filtered onto a 0.22 µm PVDF membrane (Millipore Sigma, Burlington, USA) as described(17). The filters with bacteria were placed onto 7H10-ADN plates and incubated for approximately 5 doubling times at 37°C (the following doubling times were adhered to: *M. smegmatis*: 4h; *M. abscessus*: 4h; *M. kansasii*: 17h; *M. bovis* BCG: 24h; and *M. tuberculosis*: 24h). The filters were then transferred onto inverted caps filled with 7H9-ADN without tyloxapol, glued onto 6-well plates(17). The 7H9-ADN was modified for each specific stress condition as follows: to induce acid stress, 7H9-ADN was adjusted to a pH of 5.5 with hydrochloric acid (HCl); for acid-nitrosative stress, 7H9-ADN at pH 5.5 was supplemented with 1 mM sodium nitrite (NaNO_2_); for oxidative stress, 7H9-ADN was supplemented with 5 mM hydrogen peroxide (H₂O₂); for hypoxia, well plates were placed in GasPak™ EZ anaerobic generation bags (BD); and for the multi-stress condition, 7H9-ADN at pH 5.5 was supplemented with 1 mM NaNO_2_ and 5 mM H_2_O_2_ and the 6-well plates were placed in GasPak™ EZ anaerobic generation bags (BD)(18). Next, the well plates were incubated at 37°C for 5 doubling times after which the filters were harvested for analysis. Filters were harvested by inserting them into 2 mL screw cap vials containing 0.1 mm zirconia beads and 1 mL ice-cold quenching buffer (20% (v/v) MilliQ water, 40% (v/v) acetonitrile and 40% (v/v) methanol). Cells were lysed by bead beating with a BeadBug™ 6 bead homogenizer (Benchmark Scientific, New Jersey, USA) for three cycles of 30 seconds on and 30 seconds off at 4,350 rpm, followed by a cooling step on ice, and then another round of three cycles of 30 seconds on and 30 seconds off at 4,350 rpm. After lysis, the supernatant was clarified by centrifuging for 10 min at 20,000 g at 4°C. The supernatant was then transferred to MS vials for LC-MS analysis. Media samples for LC-MS analysis were collected after the filters were harvested, and diluted 5-fold in acetonitrile:methanol (50:50, v:v). The mixture was centrifuged for 10 min at 20,000 g at 4°C after which the supernatant was transferred to autosampler vials for LC-MS analysis.

### LC-MS untargeted metabolomics

LC-MS metabolomics analyses were performed using an Agilent 1290 Infinity II LC System coupled to an Agilent 6546 Quadrupole Time of Flight MS with a Dual AJS ESI ion source (Agilent Technologies, Santa Clara, CA, USA). The ESI was operated using the following setting: gas temperature: 320 °C, drying gas: 8 L/min, nebulizer: 25 psi, sheath gas temperature: 320 °C, sheath gas flow: 11 L/min, VCap: 3,500 V, fragmentor 125 V, skimmer 50 V. The LC was fitted with a with a Cogent Diamond Hydride Type C column (150 mm × 2.1 mm, 100A, 4 μm; Microsolv) operated at 25 °C with a flow rate of 0.4 mL/min and an injection volume of 2 μL. Chromatographic separation was achieved using a gradient of buffer A (0.2% v/v formic acid in MilliQ water) and buffer B (0.2% v/v formic acid in acetonitrile): 0-2 min: 85% B, 3-5 min: 80% B, 6-7 min: 75% B, 8-9 min: 70% B, 10-11 min: 50% B, 11-14 min: 20% B, 14-24 min: 5% B, followed by re-equilibration for 10 min at 85% B(19). For untargeted metabolomic analysis, samples were analyzed in positive and negative ionization mode. For the detection of GABA-trehalose, sample were only analyzed in the positive ionization mode. Spectra were obtained in centroid mode, with an m/z range of 50-1,200 and a scan rate of 1 spectrum/s.

For data-dependent acquisition (DDA), the method above was used in combination with the Auto MS/MS functionality, at a scan rate of 4 (MS) or 2 (MS2) spectra/s. Precursor ions were selected using an absolute threshold of 10,000 counts and relative threshold of 0.01%. Abundance-dependent accumulation was activated with a target of 50,000 counts/spectrum, rejecting precursors that cannot reach the target signal within time limit. Selected precursors were actively excluded for 2 minutes after a spectrum. To reduce precursor selection of tyloxapol peaks carried over from liquid 7H9 precultures, we excluded the most tyloxapol-rich time- and mass-segments (included were 0.5-1.5 min: m/z 100-500; 1.5-2 min: m/z 100-550; 2-2.5 min: m/z 100-650; 2.5-5: m/z 100-700; 5-end: m/z 100-750). In addition to this segmented exclusion, a list of known contaminants was excluded throughout the run. Selected precursors were isolated with a narrow range (∼1.3 Da) and fragmented at a collision energy of 20 V.

### LC-MS data analysis

Raw MS data from was analyzed and processed by XCMS (version 3.12.0)(20). The IPO R package was used to optimize parameters for peak identification by the CentWave algorithm(21). Parameters deviating from default settings are: *ppm = 30; peakwidth = c(5, 65); snthresh = 6; prefilter = c(3, 2500); integrate = 2L; mzdiff = 0.00351; noise = 5000*. Peak identification was followed by retention time alignment using the ObiWarp algorithm (default settings), peak grouping (*bw = 5; minFraction = 0.8; minSamples = 1; binSize = 0.025; maxFeatures = 100*), and filling in missing peaks across samples. The resulting dataset was extensively filtered to obtain a noise-free monoisotopic feature dataset. Features were first filtered based on retention time (90-900 seconds) and m/z ratios (80–750). The removal of poor-quality peaks and the separation of overlapping peaks were carried out by manually inspecting each individual feature. Retention time and peak area were adjusted accordingly. Isotopes and other artifacts were removed with a custom-built script that utilized the CAMERA package for isotope identification(22). The final noise-free monoisotopic dataset was log2 transformed for normalization and mean-centered prior to principal component analysis (PCA) to assess the variance between samples and sample groups. For statistical comparison of feature abundance between the non-stressed control group and the stressed sample groups, we performed an unpaired t-test and calculated FDR-adjusted p-values using the Benjamini-Hochberg method for multiple comparisons. For targeted metabolite analysis, peak areas were manually integrated using a 20 ppm mass window, using MassHunter Qualitative Analysis Software (Agilent).

### Feature-based Molecular Networking and Spectral Library Search

The DDA data was used to create a feature-based molecular network with the Feature-Based Molecular Networking (FBMN) workflow described by Nothias et al. on GNPS(23, 24). In brief, the mass spectrometry data were first processed with XCMS3 and the results exported to GNPS for FBMN analysis(20, 22). The data were filtered by removing all MS/MS fragment ions within +/− 17 Da of the precursor m/z. MS/MS spectra were window-filtered by choosing only the top 6 fragment ions in the +/− 50 Da window throughout the spectrum. The precursor ion mass tolerance was set to 0.02 Da and the MS/MS fragment ion tolerance to 0.02 Da. A molecular network was then created where edges were filtered to have a cosine score above 0.55 and more than 5 matched peaks. Further, edges between two nodes were kept in the network if and only if each of the nodes appeared in each other’s respective top 15 most similar nodes. Finally, the maximum size of a molecular family was set to 100, and the lowest scoring edges were removed from molecular families until the molecular family size was below this threshold. The spectra in the network were then searched against GNPS spectral libraries(24, 25). The library spectra were filtered in the same manner as the input data. All matches kept between network spectra and library spectra were required to have a score above 0.7 and at least 5 matched peaks. The DEREPLICATOR was used to annotate MS/MS spectra(26). Finally, the molecular networks were visualized using Cytoscape software(27).

The mass spectrometry data used for the feature-based molecular network were deposited on public repository MassIVE under accession number MSV000093333. The molecular networking job can be publicly accessed at https://gnps.ucsd.edu/ProteoSAFe/status.jsp?task=fdabea21efeb4c6682f786726281819d.

### In silico structure prediction of m/z 428

For in silico structure prediction, MS2 fragmentation spectra were collected at collision energies of 10, 20 and 40 V, using the LC-MS method described above. In silico structure prediction was performed using CSI:Finger ID and MSNovelist in the SIRIUS software(28–30).

### Macrophage infection

Macrophage infections were performed as described in detail in the Supplemental Information. In brief, peripheral blood mononuclear cells (PBMCs) were isolated from whole blood from three healthy volunteers was obtained from Sanquin Bloodbank (Nijmegen, the Netherlands) using Ficoll-Paque density gradient centrifugation and resuspension in Dutch-modified RPMI 1640 medium (Invitrogen). PBMCs were seeded in Petri dishes (Corning) and incubated at 37°C in 5% CO_2_ for 2 hours after which non-adherent lymphocytes were removed and differentiation medium was added (complete RPMI supplemented with 10% (v/v) human pooled serum and 5 ng/mL macrophage colony-stimulating factor (M-CSF, Miltenyi Biotec). After incubating for 7 days, the cells were washed with PBS and harvested by adding Versene (Gibco). The cell suspensions were centrifuged at 1700 rpm for 10 min at room temperature and resuspended in complete RPMI. To activate the cells, 1 mL of the cell suspensions were diluted into 1 mL antibiotic-free medium with or without supplementation of IFN-γ, followed by incubation overnight. On day 8, the human monocyte-derived macrophages (hMDMs) were infected with *M. tuberculosis* H37Rv. To achieve this, Mtb cells were harvested by centrifuging a liquid culture and resuspended in 1 mL 0.05% (v/v) Tween 20 in PBS. Then, medium was removed from the hMDM-containing wells and the Mtb suspension was added, followed by incubation. After 4 hours, medium was removed, and cells were washed. To extract the metabolites, 1 mL of quenching buffer was added to each well, except for the hMDM-free control. Next, the suspension was bead-beaten as described above. The bacteria in the medium of the hMDM-free control were centrifuged resuspended in quenching buffer, and then subjected to bead-beating as the hMDM-containing samples. Finally, the metabolite extracts were filter-sterilized and stored at −20°C until metabolomics analysis.

### Isolation of m/z 428

A total of 120 filters with *M. tuberculosis* mc^2^ 6206 were exposed to multi-stress for 5 days and harvested as described above. The pooled lysate was dried in a SpeedVac concentrator (Savant ISS110, Thermo), resuspended in 5% (v/v) methanol in MilliQ water, and centrifuged at 20,000 g for 10 min. The clear supernatant was fractionated with an Agilent 1100 HPLC system with a diode array detector (detecting wavelengths: 210 nm, 230 nm and 250 nm) and fraction collector (G1364A), fitted with a LiChrospher 100 RP-18 endcapped column (Merck, 250 mm x 4.6 mm, 5 μm particle size) operated at 1 mL/min. After equilibration in 5% (v/v) methanol in MilliQ water, 100 μL clarified metabolite extract was injected. Two min after injection, a linear gradient was applied from 5% to 100% methanol over 30 min, followed by a gradient back to 5% methanol over 5 min. During the run, 1 mL fractions were collected. This fractionation was repeated 7 times to reach a cumulative injection volume of 700 μL. To identify fractions containing m/z 428, each fraction was analyzed by LC-MS as described above. Fractions containing the highest m/z 428 abundance were pooled, dried by rotor evaporation and stored inside a desiccator until further analysis.

### NMR spectroscopy of m/z 428

All NMR spectroscopic experiments were performed on a Bruker AVANCE III 500 MHz equipped with a Prodigy probe. Separate samples containing isolated m/z 428, α, α-trehalose, GABA, synthetic 6’-GABA-glucose or 6’-GABA-glucose were made by dissolving these compounds in D_2_O containing DSS-*d*_6_, which was used an internal chemical shift reference (^1^H: 0.00 ppm). ^1^H 1D experiments were performed with either 8 or 32 scans and a relaxation delay of 4 s. 1D selective TOCSY experiments were performed with 150 ms of spinlock mix time and 1,024 scans. 2D COSY experiments were acquired with 8 scans per increment, 256 increments in the indirect dimension and 2,048 points in the direct dimension.

### Synthesis of GABA-trehalose, GABA-glucose, and ^13^C_2_-GABA

6’-GABA-trehalose, 6’-GABA-glucose, and ^13^C_2_-GABA were synthesized as described in the Supplementary Methods. In brief, 6’-GABA-trehalose was synthesized by adding a solution of trehalose in dimethylformamide (DMF) to a solution of *N*-CbzGABA and hexafluorophosphate azabenzotriazole tetramethyl uronium (HATU) while stirring. The resulting 6’-NCbzGABA was hydrogenated using a Pd/C catalyst to yield the final product. 6’-GABA-glucose was synthesized by addition of HATU to a solution of *N*-BocGABA and glucose in pyridine. Final deprotection with TFA delivered the desired 6’-GABA-glucose. ^13^C_2_-GABA was synthesized by adding solid NaH to a solution of ^13^C_2_-triethyl phosphonoacetate in tetrahydrofuran (THF) to synthesize ^13^C_2_-ethyl 4-phthalimidobutyrate, which was then dissolved in 3M HCl and heated to a reflux to yield the final product.

### GABA-trehalose quantification in stressed Mtb

To prepare cultures for multi-stress exposure, *M. tuberculosis* mc^2^ 6206 cultures were grown to an OD_600_ of approximately 1. Cultures were then placed on ice for 15 min and centrifuged at 3216 g and 4°C for 20 min. Supernatant was discarded and cells were washed once by resuspending in 7H9-ADN medium without tyloxapol, and then centrifuging again at 3216 g and 4°C for 20 min. For the t0 hours timepoint, the cell pellet was then resuspended in 1 mL ice-cold quenching buffer and further processed and analyzed with LC-MS, as described above. Multi-stress was applied by resuspending the cell pellets in tyloxapol-free 7H9-ADN modified for multi-stress exposure, as described above, and placing the cultures in GasPak™ EZ anaerobic generation bags followed by incubation at 37°C. After 4 and 24 hours, multi-stress exposed cultures were harvested by centrifuging at 3216 g and 4°C for 20 min, resuspending the pellets in 1 mL ice-cold quenching buffer. The samples were processed and analyzed with LC-MS, as described above, but with calibration standards. To arrive at a conservative estimate of the intracellular GABA-trehalose concentration, a high estimate of the cellular volume of *M. tuberculosis* was used (8.4X10^−15^L)(31), even though lower estimates of the *M. tuberculosis* cellular volume exist (2.93X10^−16^L)(32) that would result in higher estimates of the intracellular GABA-trehalose concentration.

### GABA-trehalose synthetase activity assay

Crude *M. tuberculosis* protein lysate for the activity assay was prepared by growing a 1 L liquid *M. tuberculosis* mc^2^ 6206 culture to OD_600_ 1.0 as described above. After placing the cultures on ice for 10 min, the cells were centrifuged at 3216 g for 10 min and resuspended in a final volume of 25 mL ice-cold lysis buffer (25 mM HEPES, 75 mM NaCl, pH 7.5). Crude protein lysate was prepared by bead-beating the cell suspension as described above. Finally, the soluble protein fraction was prepared by centrifuging the crude lysate for 20 min at 20,000 g and filtering the supernatant through a 0.22μm syringe filter. All centrifugation steps were performed at 4°C.

GABA-trehalose synthetase assays were carried out by adding 27 μL crude protein lysate, soluble fraction, or (partially) purified protein to 3 μL of (10x) assay mix (consisting of 100 mM trehalose, 100 mM GABA, 50 mM ATP and 50 mM MgCl_2_). The reaction mixture was then incubated at 37°C and mixed at 600 rpm in a Thermomixer (Eppendorf). After 1 hour, the reaction was quenched by taking 20 μL of the reaction mixture and diluting it into 80 μL of 50% (v/v) methanol in acetonitrile. The quenched reaction mixture was centrifuged at 20,000 *g* for 10 min and analyzed with LC-MS as described above. Specific activity rates were determined by dividing the GABA-trehalose peak area, measured by LC-MS, by the number of minutes the reaction was incubated and dividing by protein concentration in milligram. To determine specific activity rates, protein concentrations were measured with a NanoDrop 1000 Spectrophotometer (VWR International, Radnor, USA), operated by ND-1000 v3.8.1, Proteins A280.

### GABA-trehalose synthetase purification

Soluble protein fraction was prepared as described above, but using hydroxyapatite (HA) buffer A (20 mM potassium phosphate, 100 mM NaCl, pH 7.0) instead of lysis buffer. Protein purifications were performed on an ÄKTA purifier (Cytiva, Danaher, Washington D.C., USA) with a Frac-900 fraction collector, operated at 4 °C. Following column equilibration with HA buffer A, 5 mL of *M. tuberculosis* mc^2^ 6206 soluble fraction (4 mg/mL in HA buffer A) was loaded onto a ceramic hydroxyapatite (HA) column (CHT type II, 40 μm particle size (Bio-Rad, Hercules, California), packed in an Omnifit column (Thermo Scientific, Waltham, USA) with 1 cm diameter and a packed bed volume of 8 mL), operated at 1.5 mL/min. The flowthrough (38.5 mL) was manually collected until the UV signal stabilized at close to baseline level, after which a linear gradient was started that ran from 0% to 50% buffer B (1 M potassium phosphate, 100 mM NaCl, pH 7.0) over 20 column volumes. Fraction (10 mL) collection was started using the Frac_900 function as soon as a stable increase of ∼5 mAU was observed. The resulting fractions were concentrated approximately 10-fold with a Vivaspin 20 10kDa MWCO centrifugal concentrator (Sartorius, Göttingen, Germany), and finally buffer-exchanged to 15Q buffer A (50mM Tris, 100mM NaCl, pH 8.0) with PD midiTrap G-25 cartridges (Cytiva) using the gravity protocol at 4 °C. The resulting concentrated fraction in 15Q buffer A were then subjected to the GABA-trehalose synthetase assay (see above) or purified further. The fractions exhibiting the highest specific GABA-trehalose synthetase activity (i.e., fraction 1, which eluted between 16.3 mS/cm and 21.8 mS/cm) of two separate HA purifications were pooled for further fractionation. The second fractionation step was performed with a pre-packed Source 15Q 4.6/100 PE column (Cytiva) operated at 0.5 mL/min. After the column was equilibrated in 15Q buffer A, the pooled active fractions from the HA fractionation (5 mL at 1.78 mg/mL) were loaded. After washing with 20 mL buffer A, a linear gradient was started from 0% to 65% 15Q buffer B (50 mM Tris, 1M NaCl, pH 8.0) over 20 column volumes. Fractionation collection was done as for the HA purification, but with fractions of 2 mL and as soon as a stable increase of ∼1 mAU was observed. The resulting fractions were concentrated, buffer-exchanged to 15Q buffer A and assayed for GABA-trehalose synthetase activity as described for the HA fractions. The three factions exhibiting the highest GABA-trehalose synthetase activity, i.e., fractions 5, 6, and 7 (eluting between a conductivity of 24.38 mS/cm and 77.66 mS/cm) were selected for proteomic analysis.

### Proteomics

The selected protein fractions were diluted 1:1 (v/v) with 8 M urea, 10 mM Tris, pH 8.0. Reduction was achieved by adding 1 μL of 10 mM dithiothreitol per 50 μg of protein, followed by a 30 min incubation at room temperature. Reduced proteins were then alkylated by adding 1 μL of 50 mM 2-chloroacetamide with 50 mM ammonium bicarbonate per 50 μg of protein and incubating for 20 min at room temperature in the dark. Proteins were then digested by adding 1 μg of trypsin per 50 μg of protein and incubating overnight at 37°C. Peptides were then diluted 1:1 (v/v) with 2% (v/v) trifluoroacetic acid (TFA) and concentrated in a SpeedVac concentrator (Savant ISS110, Thermo), until no visible liquid was left. Peptides were then dissolved in 20 μL 0.1% TFA and stored at −20°C until proteomic analysis. Proteomics analyses were performed using an Evosep One nanoflow liquid chromatography system (Evosep) connected online via a CaptiveSprayer electrospray ionization source to a timsTOF Pro2 mass spectrometer (Bruker Daltonics). For each sample, 200 ng of tryptic peptides were injected and separated using the 30 samples-per-day Evosep method on a 30SPD endurance column (EV1106: 150 x 0.15 mm, ReproSil-Pur C18, 1.9 µm particles; Evosep) at 45 °C. Mass spectrometry data acquisition was performed in positive ionization mode using the default 1.1 s duty cycle dda-PASEF instrument method for proteomics (0.6-1.6 1/K0 mobility range, 100-1,700 *m/z* mass range, 10 PASEF frames, dynamic exclusion enabled for 30 s)(33). Peptide identification and protein inference was performed in real-time using the ProLuCID search engine(34) in PaSER v2023 (Bruker Daltonics) with the following settings: protein sequence database: UniProt Mycobacterium tuberculosis strain ATCC 25618 / H37Rv OX=83332 date=2022-10-20, tryptic specificity, 20 ppm precursor and fragment ion tolerance, carbamidomethyl (Cys) as fixed modificiation, Oxidation (Met) as variable modification, dta_select 1% protein FDR threshold, TIMScore enabled. Protein label-free quantitation was performed using the PaSER dda-MBR module at 10 ppm mass tolerance, 0.03 1/K0 mobility tolerance, 30 s retention time tolerance, minimum XIC traces correlation of 0.7, and maximum RMSE isotopic ratio 0.2.

### Heterologous expression of Rv1722, Rv2411c, and Rv2567

Linear fragments encoding Rv1722, Rv2411c, and Rv2567 (Uniprot accessions P71980, P9WLA9 and P9WL97, respectively) were modified with a C-terminal GS-linker (1x GGGGS), codon optimized, synthesized, and assembled into a pET28(a) vector by Twist Bioscience (South San Francisco, USA) so that the C-terminal GS linker was followed by a 6xHis-tag. The resulting sequence-verified plasmids (50 ng) were used to transform chemically competent *E. coli* BL21DE3 (Thermo Scientific) using heat shocking at 42°C for 1 min. Following heats shock, 700 μL of LB medium was added and cells were allowed to recover at 37°C for 1 hour. The bacteria were then centrifuged at 2,500g for 2 min, resuspended in 100 μL LB medium, spread onto an LB agar plate containing 50 μg/mL kanamycin sulfate, and incubated overnight at 37°C. For each gene, a colony was picked from the overnight plates and used to inoculate TB medium (2% (w/v) peptone, 2.4% (w/v) yeast extract, 0.2313% (w/v) KH_2_PO_4_, 1.6433% (w/v) K_2_HPO_4_⋅ 3 H_2_O, 0.4% (v/v) glycerol) supplemented with 50 μg/mL kanamycin sulfate. Next, the cultures were incubated at 37°C and 120 RPM until the OD_600_ was between ∼0.6 and 1. Then, protein production was induced by adding 500 nM IPTG (final concentration) and cultures were incubated overnight at 25°C. After the overnight incubation, the cultures were placed on wet ice for ∼10 min, centrifuged at 5,000 g and 4°C for 10 min, and resuspended in 50 mM potassium phosphate + 50 mM NaCl, pH 7.0. Finally, crude lysate was prepared by lysing the resuspended cells with a Sonopuls GM3200 with KE-76 attachment (Bandelin, Berlin, Germany), through three one-minute rounds of 1 s on and 1 s off sonication at 55% amplitude on ice.

### Rv1722 purification

For HisTrap purification of Rv1722, a 2 L expression culture that was cultured as described above was harvested by placing the culture flasks on wet ice for ∼15 min and centrifuging at 7,000 g and 4 °C for 10 min. Cells were resuspended in HisTrap buffer A (50 mM HEPES, 500 mM NaCl, 10 mM imidazole, pH 7.5) to a final volume of ∼80 mL and lysed using sonication as described above. The lysate was then ultra-centrifuged at 120,000g and 4 °C for 30 min (Fixed angle Ti45 Rotor, Optima XE90, Beckman Coulter). The supernatant was filtered through a syringe filter (0.22 μm pore size) to obtain the soluble fraction. The purification was done with a 5 mL HisTrap HP (Cytiva, Danaher, Washington D.C., USA) attached to an ÄKTA purifier (Cytiva, Danaher, Washington D.C., USA), operated at 4 °C and 1 mL/min flowrate. The soluble fraction was loaded at 1 mL/min onto the HisTrap column after it was equilibrated in HisTrap buffer A. After the flow through had passed and the UV absorbance stabilized, the column was washed with 49.2 mM imidazole by applying a step gradient to 8% HisTrap buffer B (50 mM HEPES, 500 mM NaCl, 10 mM imidazole, pH 7.5). After the following UV absorbance peak had passed and UV signal was back at approximately baseline level and stabilized, a step gradient to 100% HisTrap buffer B was initiated. A single 3 mL fraction was manually collected after the UV absorbance increased by ∼450 mAU. The collected fraction was then concentrated to a volume of 1 mL and buffer exchanged to 50 mM HEPES, 200 mM NaCl, 5 mM MgCl_2_, pH 7.5, as described under ‘GABA-trehalose synthetase purification’ and stored on ice until use for a GABA-trehalose synthetase activity assay.

### GABA-trehalose formation upon GABA exposure

*M. tuberculosis* mc^2^ 6206-laden filters were prepared as described under ‘Mycobacterial stress exposure’ and then transferred onto inverted caps filled with 7H9-ADN liquid medium containing 0-100 µM GABA. After 24 h of incubation at 37 °C, filters were harvested as described under ‘Mycobacterial stress exposure’ and analyzed by LC-MS untargeted metabolomics in positive mode, as described under ‘LC-MS untargeted metabolomics’. For the labelling experiment, 1 mM ^12^C-GABA or ^13^C_2_-GABA was added to the culture media as described above. The labeling of GABA-trehalose was assessed using Profinder 10.0 (Agilent Technologies, Santa Clara, CA, USA).

### Phylogenetic analysis of Rv1722

To identify putative homologs of Rv1722 in publicly available *Mycobacterium* genomes, the NCBI datasets utility was used to download the proteome sequences of all *Mycobacterium* records in GenBank that contained annotation data (6,966 on 13 February 2025). Blastp was used with the Rv1722 amino acid sequence as query against each proteome. The e-value threshold was set at 10^−3^ and the minimum percentage identity to 50%. *Mycobacterium* species with hits were collected in supplementary table 1. Annotations for slow- and fast-growing species were added if a traceable author statement was available (TAS). The PubMed links in the table refer to these TASs.

Data to generate the phylogenetic tree was retrieved from a custom database of assembled genome sequences of 227 NTM type strains (version 1.3.1, https://github.com/Bioinf-MMB-Rumc/MyCodentifier), with full genome sequences retrieved from the NCBI genome database(35). The phylogenetic tree was built using mashtree (version 1.2.0)(36), with 100 bootstrap replicates and with *Hoyosella altamirensis* included as an outgroup (based on Tortoloni et al.(37)). Growth rates were derived from Tortolini et al.(37), while novel species were classified using species determination references available through NCBI. The DNA sequence of *rv1722* from the complete genome of *Mycobacterium tuberculosis H37rv*(38) was used as a reference sequence to screen for its presence in the type strains. *rv1722* was identified and extracted from each type strain using NCBI Nucleotide BLAST(39), applying a cutoff threshold of 50% sequence similarity. Visualization of the phylogenetic tree was performed in R using the *ggtree* package (version 3.17.1.1)(40).

### Stable-isotope labeling

*M. tuberculosis* mc^2^ 6206-laden filters were prepared as described under ‘Mycobacterial stress exposure’, transferred onto inverted caps filled with the ^13^C_2_-acetate-containing 7H9-ADN liquid medium (prepared by replacing the D-glucose and glycerol with 0.2% w/v ^13^C_2_-acetate sodium salt (99% isotopic purity, Millipore Sigma, Burlington, USA)), and then immediately exposed to hypoxia as described under ‘Mycobacterial stress exposure’ or placed in an open zip lock bag without reagent as oxic control. After 24 hours, filters were processed and analyzed by LC-MS untargeted metabolomics analysis as described above. For stable isotope tracing with D_4_-succinate as substrate, *M. tuberculosis* mc^2^ 6206-laden filters were first preincubated on non-labeled succinate-containing 7H9-ADN liquid medium (prepared by replacing the D-glucose and glycerol with 30 mM succinic acid and setting the pH to 6.6 using 6 M NaOH). After 24 hours of preincubation the filters were transferred to inverted caps filled with D_4_-succinate-containing multi-stress 7H9-ADN liquid medium (prepared by replacing the D-glucose and glycerol with 30 mM D_4_-succinic acid (Succinic acid-2,2,3,3-d4, 98% isotopic purity, Millipore Sigma, Burlington, USA), supplementing with 1 mM NaNO_2_ and 5 mM H_2_O_2_, setting the pH to 5.5 using 1 M NaOH), and immediately exposing to hypoxia as described under ‘Mycobacterial stress exposure’. For the non-stressed control, filters were transferred onto inverted caps filled with 7H9-ADN liquid medium that was prepared by replacing the D-glucose and glycerol with 30 mM D_4_-succinic acid and setting the pH to 6.6 using 6 M NaOH. After 24 hours filters were processed and analyzed by LC-MS untargeted metabolomics analysis as described above. Isotopologs of target metabolites were extracted using Profinder 10.0 (Agilent Technologies, Santa Clara, CA, USA). Aspartate, glutamate, GABA, and GABA-trehalose were analysed in the positive ionization mode, and (iso-)citrate, succinate, fumarate, and malate in the negative ionization mode.

## Results

### Phagolysosome-mimicking stresses induce an unknown metabolite cluster in *Mycobacterium tuberculosis*

To discover unknown metabolites involved in mycobacterial stress response, we first screened for stress-responsive metabolites. We exposed the BSL-2 level auxotrophic Mtb strain mc^2^ 6206 to stresses that mimic conditions of the phagolysosome and profiled metabolic changes using untargeted metabolomics (Fig 1A). The resulting metabolite profiles show that stress-induced changes are stress-specific and vary in quality (Fig 1B) as well as quantity (Fig 1C-G). Hypoxia, acid-nitrosative and multi-stress metabolomes were most impacted. Although most features (LC-MS peaks) were present in both stressed mycobacteria and non-stressed controls, several hundred features were unique to stressed mycobacteria (Fig 1C-G).

**Figure 1.**
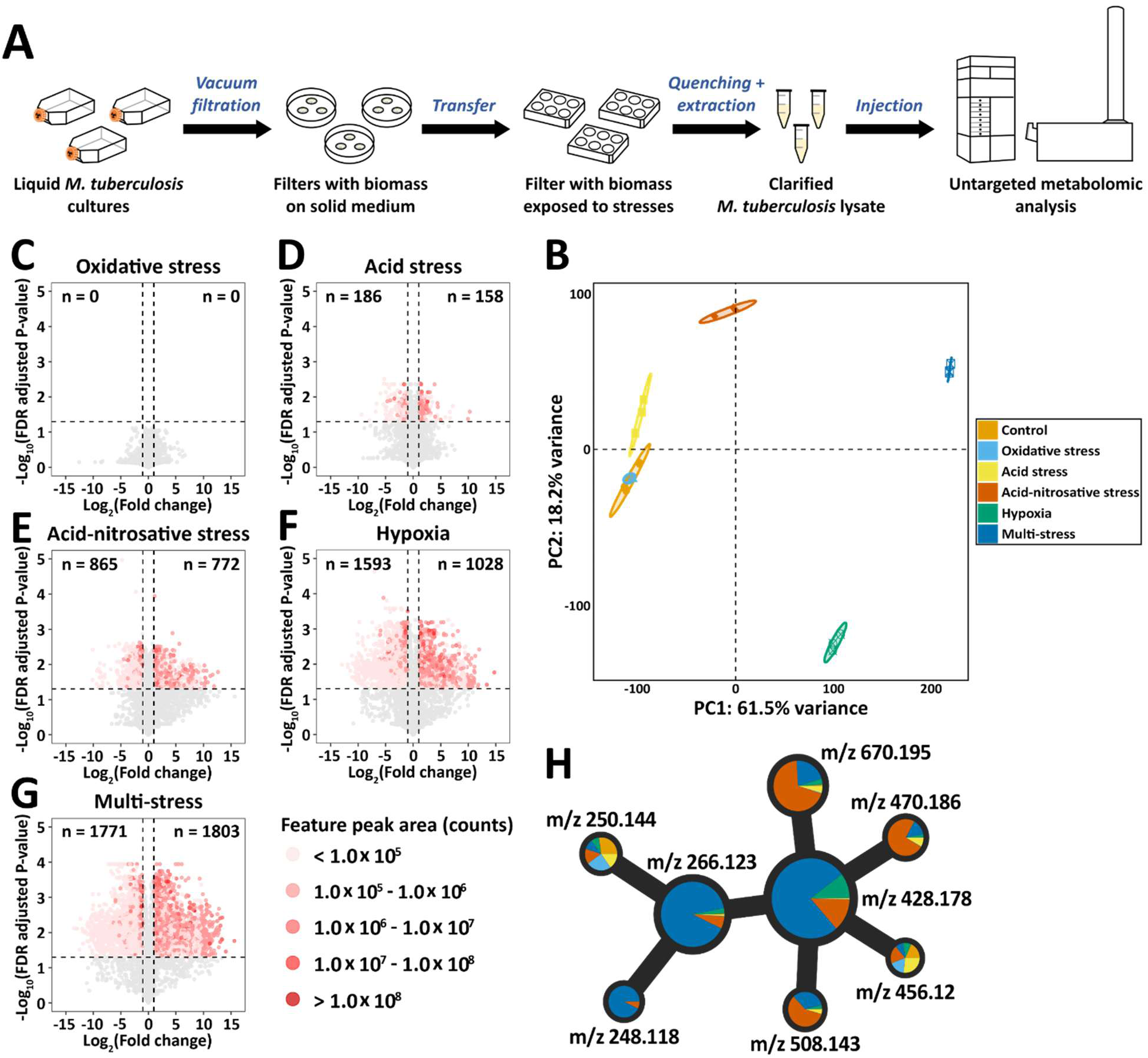
Untargeted metabolomics reveals stress-specific metabolic changes in Mtb. **A)** Graphical representation of the experimental setup. Mtb was grown in liquid 7H9 broth and then transferred to filters on 7H10 agar. After 5 days, filters were placed on top of a reservoir containing liquid media mimicking phagolysosome stresses. Samples for LC-MS analysis were taken by quenching the filters in ice-cold acetonitrile-methanol-water, followed by bead-beating. **B)** PCA plot visualizing the metabolomes of Mtb exposed to phagolysosome-mimicking stresses; Control (no stress applied), oxidative stress (5 mM H2O2), acid stress (pH 5.5), acid-nitrosative stress (1 mM NaNO2, pH 5.5), hypoxia (∼1% O2), and multi-stress (all stress conditions combined). Replicates are plotted individually (n=3). **C-G)** Volcano plots showing stress-induced metabolome changes. Each data point represents the average log2 transformed fold change and -log10 transformed FDR-adjusted *P*-value of a feature (stress condition versus control, n=3). Features with a *P*-value <0.05 and absolute log2(FC) >1 are colored by the maximum feature intensity in log10(counts). The total number of features in each quadrant is shown in the corner of each quadrant. **H)** Subnetwork of a feature-based molecular network (Fig S1) that shows a cluster of highly stress-responsive but unidentified features. Each node represents a feature and is depicted as a pie chart that represents the mean relative peak area across stresses (n=3). The colors indicate the applied stresses and match the colors in figure 1B. Node sizes represent the summed feature intensity across all conditions on a log10 scale. Nodes are connected via edges if the MS^2^-spectra are similar (Cosine score >0.55 and more than 5 matched peaks). To generate this subnetwork, the original subnetwork (Fig S1) was manually curated by improving peak integration and removing nodes representing erroneous features or isotopes.

To further explore these stress-responsive features, we generated a feature-based molecular network which visualizes chemical similarity of known and unknown metabolites in a network (Fig S1)(23). Most features in our molecular network were not unique to stress. However, members of one cluster of unknown but structurally related metabolites were highly responsive to stress (Fig 1H). This cluster of 8 features was centered around two features with molecular weights of m/z 428.178 [M+H]^+^ and 266.124 [M+H]^+^. These two features accounted for most of the cluster’s total signal intensity (85% and 12%, respectively) and were also among the highest intensity features of stressed Mtb metabolomes in general. Notwithstanding their abundance, extensive database searches(25, 41–43) and in silico structure prediction based on high-resolution MS and MS^2^ spectra(28, 29), failed to yield plausible chemical structures for m/z 428 and m/z 266. Taken together, our metabolomic screen revealed two highly abundant but unknown stress-responsive metabolites in Mtb.

### Unknown metabolites are rapidly produced by pathogenic mycobacteria exposed to infection conditions

Before proceeding with structural elucidation of the unknown metabolites m/z 428 and 266, we first aimed to explore their distribution across pathogenic and non-pathogenic mycobacteria, and verify their presence under infection conditions.

We determined the formation of m/z 428 and m/z 266 in non-pathogenic *M. smegmatis*, the opportunistic pathogens *M. abscessus* and *M. kansasii*, and the professional pathogen *M. tuberculosis* (strain mc^2^ 6206) under the phagolysosome stresses described above. Both metabolites almost exclusively accumulated in *M. tuberculosis* (Fig 2A-B). While the single stresses hypoxia and acid-nitrosative stress induced low-level production, multi-stress resulted in the highest levels. Under multi-stress conditions, only *M. tuberculosis* and *M. kansasii* produced m/z 428, but levels in *M. kansasii* were ∼20-fold lower (Fig 2A-B).

**Figure 2.**
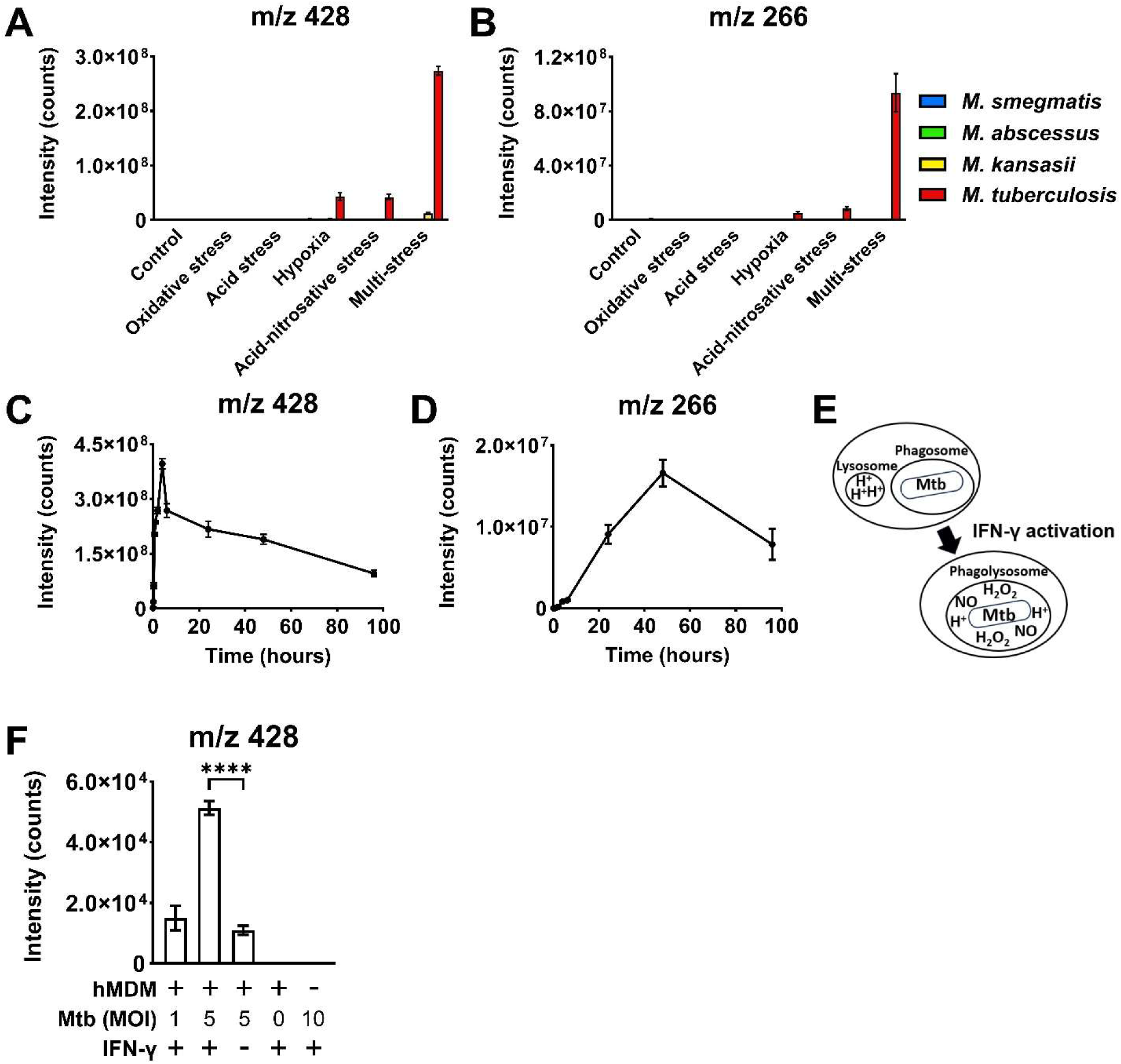
Unknown metabolites are rapidly produced by pathogenic mycobacteria exposed to infection conditions. **A&B)** Production of m/z 428 and 266 in pathogenic and non-pathogenic mycobacteria exposed to phagolysosome-like stresses. Mycobacteria were exposed to stresses for 5 doubling times using the setup described in figure 1A, followed by LC-MS determination of m/z 428 and 266 levels. **C&D)** Time course of m/z 428 and 266 formation upon stress exposure. Mtb was exposed to multi-stress without hypoxia and formation of m/z 428 and 266 was followed over time using LC-MS. **E)** Graphical representation of the IFN-γ-induced maturation of the phagosome into the phagolysosome in human macrophages. **F)** Formation of m/z 428 in Mtb H37Rv in human monocyte derived macrophages (hMDM) originating from three different donors, in the presence and absence of IFN-γ. Statistical testing was done with an unpaired two-tailed t-test. MOI: multiplicity of infection (MOI). **** = *P* <0.0001. All data are presented as mean +/− standard deviation (n=3).

To gain more insight into the kinetics of m/z 428 and 266, we exposed *M. tuberculosis* mc^2^ 6206 to multi-stress without hypoxia and monitored metabolite levels over 96 hours. Unknown m/z 428 appeared within 10 minutes after stress exposure and rapidly increased up to 4 hours (Fig 2C). Unknown m/z 266 peaked later, at 48h, and reached intensity levels that were ∼25-fold lower (Fig 2D).

Although our screens were designed to mimic phagolysosome conditions, the *in vitro* stresses and auxotrophic *M. tuberculosis* strain mc^2^ 6206 do not fully capture infection conditions. To confirm that our results are relevant under actual infection conditions, we next assessed production of the unknown metabolites in a macrophage infection model with primary human monocyte-derived macrophages (hMDMs) and wild-type, virulent *M. tuberculosis* (H37Rv) (Fig 2E). Infection of hMDMs led to clear m/z 428 production that scaled with the multiplicity of infection (Fig 2F). We did not detect m/z 266 but assume the low number of bacteria in this experiment to be the cause. Activation of the hMDMs with interferon-gamma (IFN-γ), which results in phagolysosome maturation(44), led to a 5-fold increase in m/z 428 levels, confirming its responsiveness to physiologically relevant stresses (Fig 2E&F).

Collectively, these results show that the unknown metabolites m/z 428 and 266 are rapidly formed in pathogenic mycobacteria exposed to infection conditions, showing their potential role in metabolic adaptation and pathogenesis.

### Unknown metabolites m/z 428 and 266 are GABA-trehalose and GABA-glucose

Despite their high abundance and apparent role in mycobacterial stress adaptation, m/z 428 and 266 were unknown metabolites and refractory to computational annotation. Therefore, we cultured a large amount of Mtb biomass, exposed it to the multi-stress condition and extracted the mycobacterial metabolites for purification. Reversed-phase HPLC fractionation resulted in several fractions with high m/z 428 levels that we pooled and concentrated for NMR analysis. A combination of various ^1^H and ^13^C NMR methods suggested the presence of an α,α-trehalose substructure, which was confirmed with a standard (Fig S2). The mass difference between m/z 428 and trehalose, in combination with the NMR spectra, further suggested the presence of a γ-amino-butyric acid (GABA) fragment. Together these results implied that m/z 428 is trehalose with an O6-linked GABA moiety. To confirm this putative identification, we synthesized 6’-GABA-trehalose (GABA-trehalose) and found that its LC-MS retention time (Fig 3A&B), MS^2^ fragmentation spectrum (Fig 3E&F) and NMR spectrum (Fig 3I) matched those of unknown m/z 428. These results not only established that m/z 428 is GABA-trehalose (Fig 3J) but also led to the putative identification of m/z 266 as 6’-GABA-glucose (Fig 3K), which was similarly confirmed with a synthesized standard (Fig 3C&D, G&H). Using the synthesized GABA-trehalose standard, we next quantified the intracellular concentration of GABA-trehalose upon multi-stress exposure. Based on a high estimate of bacterial cell volume, we detected a conservative intracellular concentration of approximately 10 mM, placing GABA-trehalose among the most abundant Mtb metabolites (Fig S3). The other, low-abundant molecular network cluster nodes (Fig 1H) remain unidentified. Taken together, these results thus establish GABA-trehalose and GABA-glucose as novel mycobacterial stress response metabolites.

**Figure 3.**
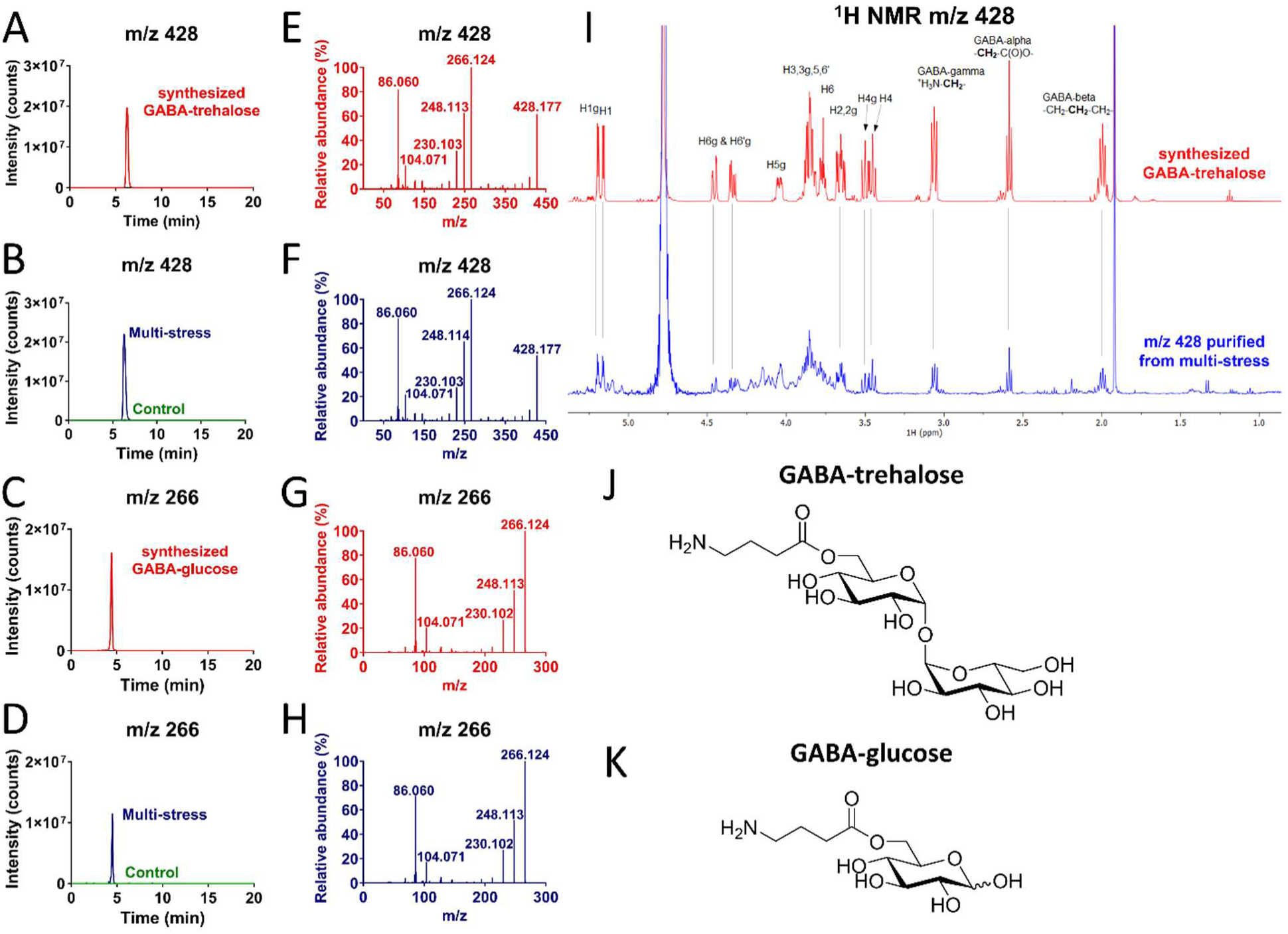
Identification of unknown metabolites m/z 428 and 266 as GABA-trehalose and GABA-glucose. **A-D)** LC-MS chromatograms of synthesized GABA-trehalose and GABA-glucose (red) compared to unknown metabolites m/z 428 and m/z 266 in multi-stress-exposed Mtb cultures (blue) and non-stressed control Mtb cultures (green). **E-H)** MS^2^ fragmentation spectra of synthesized GABA-trehalose and GABA-glucose (red) and unknowns m/z 428 and m/z 266 in multi-stress-exposed Mtb cultures (blue). **I)** Stacked ^1^NMR plots of the synthesized GABA-trehalose (red) and m/z 428 purified from multi-stress exposed Mtb cultures (blue). **J&K)** Structural formulas of GABA-trehalose and GABA-glucose.

### GABA-trehalose formation is driven by GABA accumulation

To explore potential drivers of GABA-trehalose formation, we compared the levels of GABA and trehalose to those of GABA-trehalose (as shown in Fig 2C) upon multi-stress exposure (Fig 4A). Interestingly, we found that GABA-trehalose levels were strongly correlated with GABA levels, but not with trehalose levels. We therefore argued that the rapid formation of GABA-trehalose could be a GABA-driven process. In line with this hypothesis, we found that GABA levels peaked just before GABA-trehalose (Fig 4A). To confirm that GABA-trehalose formation is a GABA-driven process, we incubated Mtb with GABA without applying stress, and found extensive, concentration-dependent, GABA-trehalose formation (Fig 4B). Moreover, exposing Mtb to ^13^C_2_-GABA led to the formation of ^13^C_2_-GABA-trehalose, showing that the produced GABA-trehalose originated from externally supplied GABA (Fig 4C). Together, these experiments suggest that the enzyme that produces GABA-trehalose is constitutively present but produces GABA-trehalose only when GABA levels increase.

**Figure 4.**
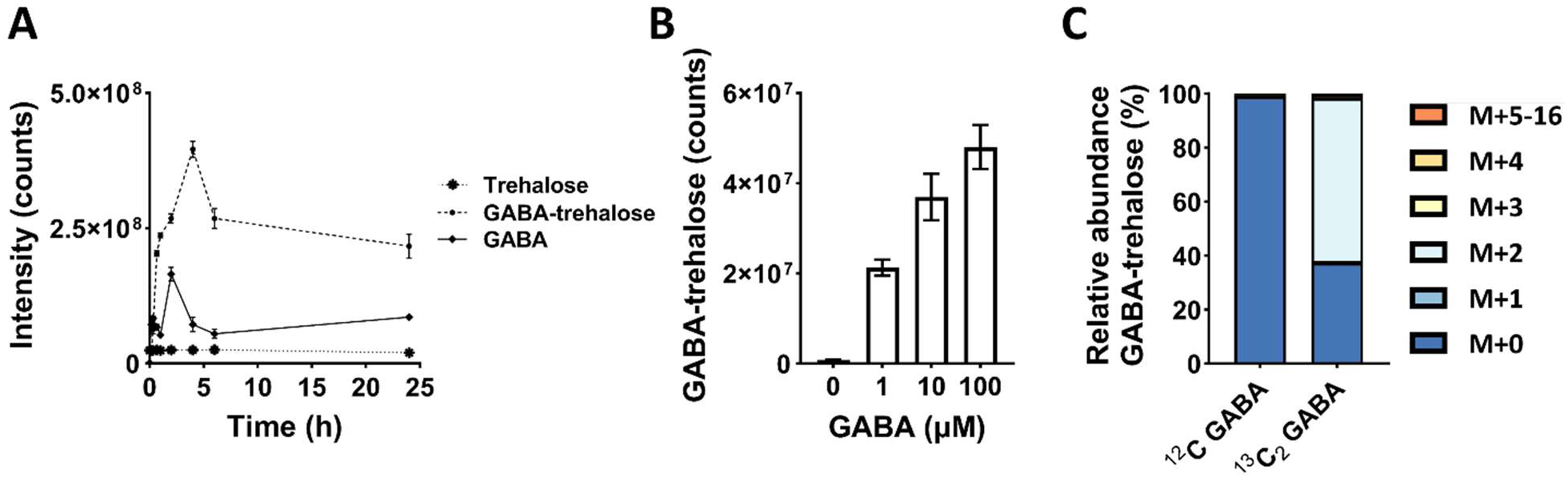
GABA-trehalose formation is substrate-driven. **A)** GABA, trehalose, and GABA-trehalose production by Mtb exposed to multi-stress conditions without hypoxia, using the setup depicted in Fig 1A; GABA-trehalose data is identical to Fig 2C. The dashed line for GABA-trehalose shows same data as was plotted in Fig 2C. Data are presented as mean +/− standard deviation (n=3). **B)** GABA-trehalose production by Mtb 24 h after exposure to the indicated GABA concentrations under non-stress culture conditions. Data are presented as mean +/− standard deviation (n=3). **C)** Isotopologs of GABA-trehalose produced by Mtb 24 hours after adding 1 mM ^12^C-GABA or ^13^C2-GABA to non-stress culture media. Values are means of replicate filters (n=3) and corrected for natural isotope abundance.

### Hypoxia blocks GABA degradation

To explore the effect of stress on flux through the TCA cycle, GABA shunt, and GABA-trehalose, we next incubated Mtb with the stable isotope-labeled carbon source ^13^C_2_-acetate under normoxia and hypoxia and assessed the labeling pattern of TCA and GABA shunt metabolites. We focused on the single stress hypoxia since it has been studied in Mtb before and is less complex to interpret than multi-stress.

In line with previous reports, hypoxia remodeled TCA cycle activity, as observed by reduced ^13^C-labeling and a strong accumulation of succinate(45) (Fig 5). The accumulation of succinate was mirrored by similar increases in GABA and GABA-trehalose. Both GABA and GABA-trehalose were extensively labeled, confirming their active role in central carbon metabolism under hypoxia. In addition to succinate accumulation, previous reports also showed succinate excretion(7, 46). To assess whether GABA and GABA-trehalose are also excreted, we next analyzed culture medium samples and found that GABA is also excreted in high amounts (Fig S4). In contrast, GABA-trehalose was only present at very low levels in the medium. These results thus show that GABA rapidly accumulates under hypoxia, and that this accumulation leads to excretion and GABA-trehalose formation.

**Figure 5.**
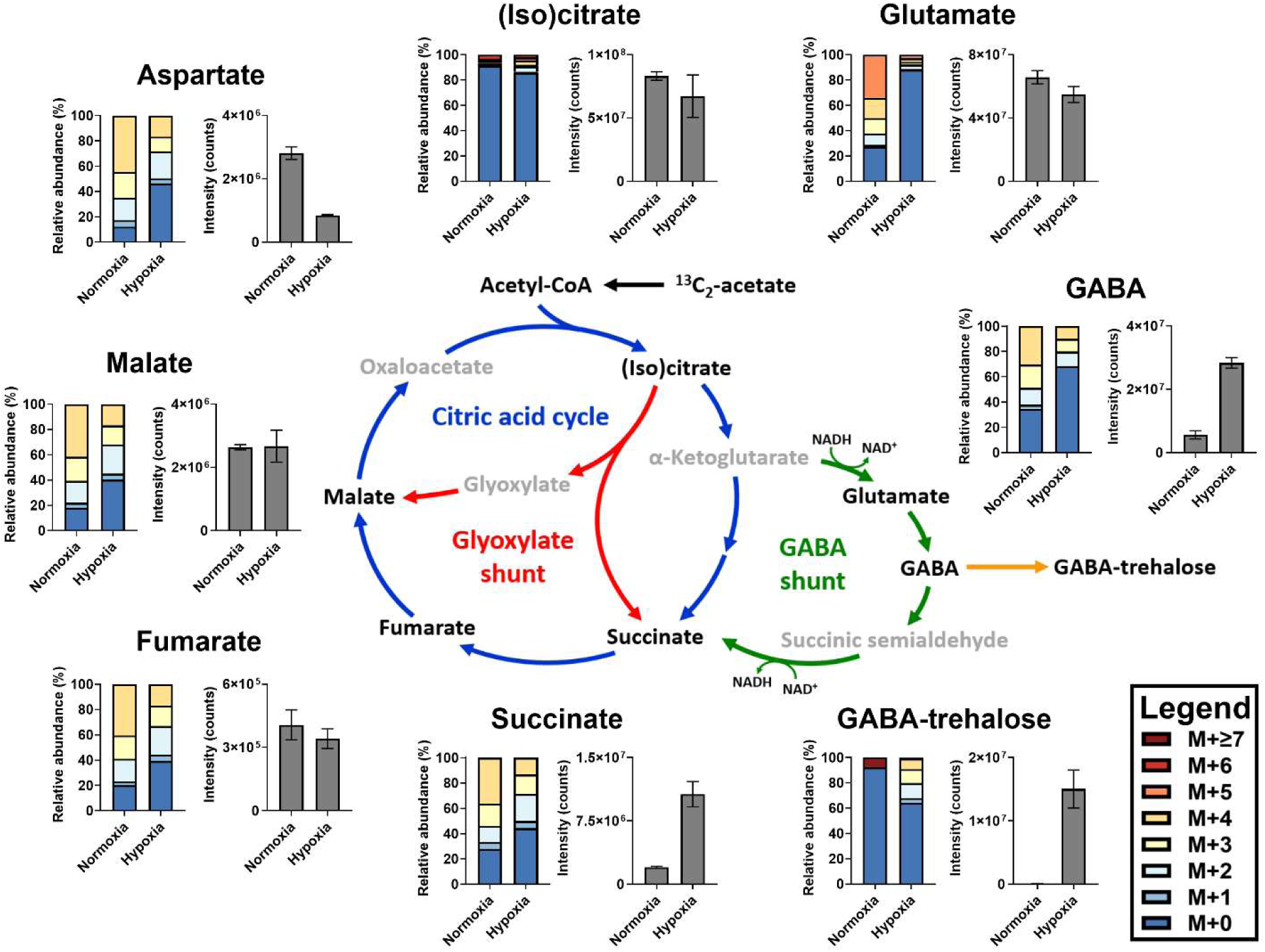
Hypoxia leads to remodeling of the TCA cycle and the GABA shunt. Relative abundances of the isotopologs and total abundances of intermediates in central carbon metabolism 24 hours after transferring in Mtb-laden filters to medium containing ^13^C2-acetate, with or without the application of hypoxia. M+n indicates the number of ^13^C atoms incorporated into the metabolite. For GABA-trehalose, the number of incorporated ^13^C atoms higher than or equal to 7 are summed. Metabolites that were not detected are depicted in gray. Data are presented as mean (n=3) and are corrected for natural isotope abundance.

GABA is an intermediate in the GABA-shunt, which runs from α-ketoglutarate to succinate, and involves two redox reactions (Fig 5)(47). Previous work on Mtb has shown that hypoxia induces succinate accumulation and strongly increases the NADH/NAD+ ratio(45, 46). On the one hand this shift in redox balance promotes the first step of the GABA-shunt (conversion of α-ketoglutarate to glutamate), but on the other hand the same shift in redox balance, along with the succinate accumulation, could inhibit the last step (succinic semialdehyde to succinate) leading to GABA accumulation.

Since the enzymes in the GABA-shunt are in principle reversible, we hypothesized that the high NADH/NAD+ ratio and succinate level under hypoxia could potentially even reverse the last steps of the GABA-shunt, producing GABA from succinate. While our labeling data supported a reversal of the final steps of the GABA shunt (Fig 5), additional stable-isotope tracing experiments with D_4_-succinate clearly showed that a reverse GABA-shunt was not active even under multi-stress conditions (Fig S5), confirming previous studies showing that succinic semialdehyde dehydrogenase is not readily reversible in vitro(48). We therefore conclude that GABA is most likely formed from glutamate via the canonical direction, and that the mismatch in glutamate and GABA labeling is caused by unlabeled glutamate present in the culture media or an unknown source of labeled GABA.

Together, these results show that hypoxic conditions lead to GABA accumulation, likely due to decreased conversion to succinate, and subsequent overflow of GABA via excretion and GABA-trehalose production.

### GABA-trehalose is produced by the uncharacterized enzyme Rv1722

GABA-trehalose and GABA-glucose have never been reported in literature and therefore represent an entry point into an uncharted metabolic pathway. To access this entry point we sought to identify the enzyme that produces GABA-trehalose. Based on chemical structure and GABA-dependence, we inferred that GABA and trehalose were likely substrates for the biosynthesis of GABA-trehalose. However, initial assays with crude Mtb protein lysate, GABA, and trehalose did not yield GABA-trehalose. Addition of suspected cosubstrates revealed that ATP was sufficient to drive the enzymatic ligation of GABA and trehalose in an Mtb crude lysate (Fig S6A). To identify the enzyme responsible for GABA-trehalose biosynthesis, we next performed activity-guided protein fractionation. Consecutive protein fractionation with hydroxyapatite (HA, Fig 6A) and strong anion-exchange chromatography (15Q, Fig 6B) resulted in an enriched fraction that we analyzed using shotgun proteomics, along with the two neighboring fractions. Although the enriched fraction contained 660 detectable proteins, filtering based on correlation with the enzyme activity profile, protein intensity, and presence of an ATP-binding domain resulted in a limited number of candidate enzymes, of which we selected Rv1722, Rv2411c and Rv2567 for further testing. Codon-optimized and His-tagged versions of Rv1722, Rv2411c and Rv2567 were expressed in *E. coli*, followed by GABA-trehalose biosynthesis tests in crude protein lysates. Only protein lysates of *E. coli* expressing Rv1722 produced GABA-trehalose, suggesting that Rv1722 catalyzes GABA-trehalose biosynthesis (Fig 6C). Activity assays with purified heterologously expressed His-tagged Rv1722 confirmed that it catalyzes GABA-trehalose synthesis (Fig 6D). In contrast to trehalose, glucose did not serve as a substrate for GABA-glucose formation and no inhibitory effect of glucose on GABA-trehalose biosynthesis could be observed (Fig S6B&C). Based on these findings, we conclude that Rv1722 catalyzes GABA-trehalose formation through the reaction shown in Fig 4E and name it GABA-trehalose synthetase (GabtS).

**Figure 6.**
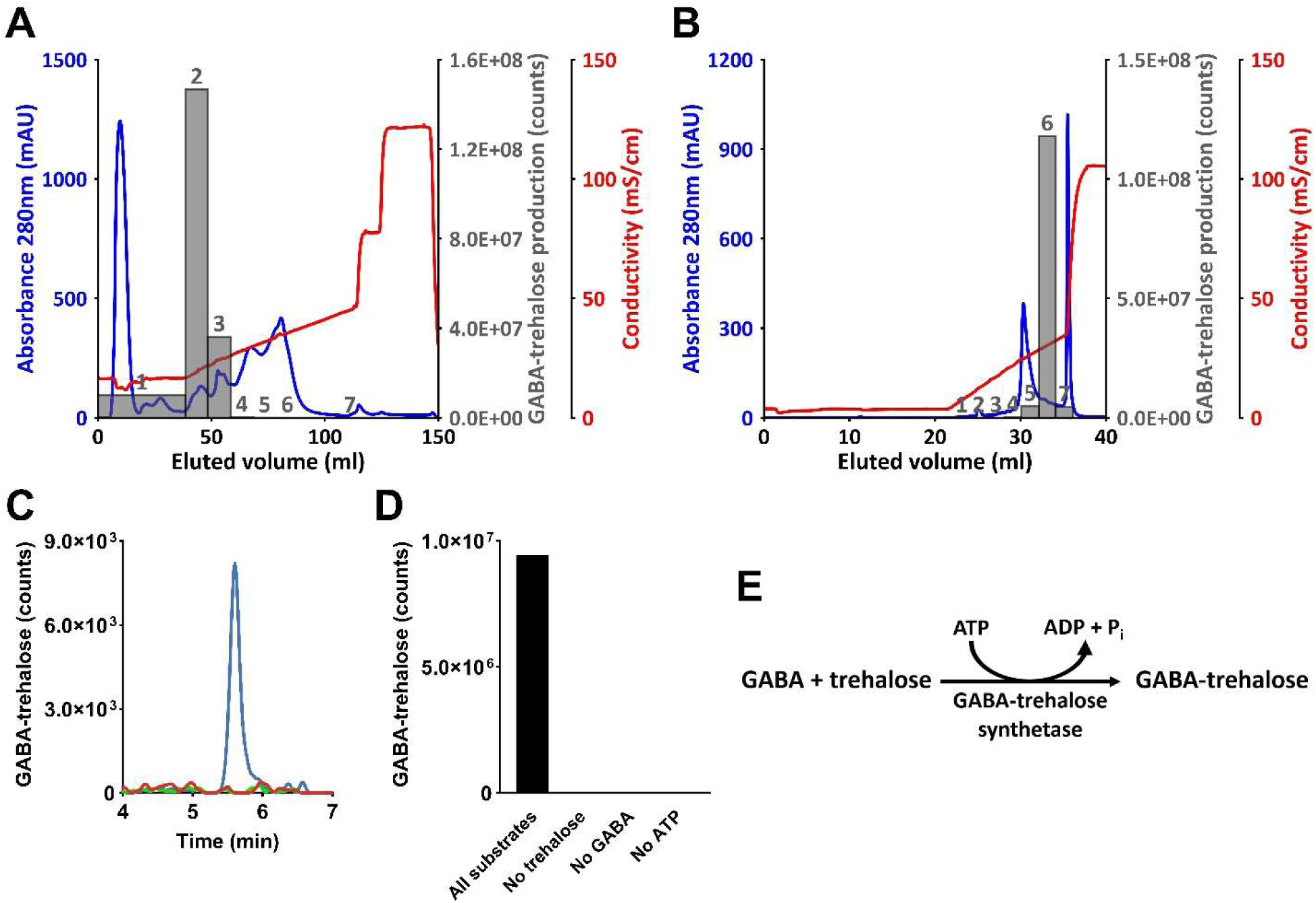
GABA-trehalose is produced by the uncharacterized enzyme Rv1722. **A)** FPLC chromatograms of activity-guided fractionation of the GABA-trehalose synthetase from crude Mtb protein lysate with hydroxyapatite chromatography. **B)** Same as A, but with 15Q chromatography. Grey numbered bars show the produced GABA-trehalose in each of the protein fractions. **C)** Superimposed LC-MS chromatograms of the GABA-trehalose production in the crude lysates of *E. coli* BL21 DE3 transformed with vectors containing constructs encoding Rv1722, Rv2411, and Rv2567. **D)** GABA-trehalose synthesis by His-trap-purified heterologously expressed Rv1722 with trehalose, GABA, and ATP as substrates, and under omission of GABA, trehalose, or ATP. **E)** Biosynthetic reaction for the Rv1722-catalyzed ATP dependent formation of GABA-trehalose from GABA and trehalose. n=1 for all experiments.

### Rv1722 homologs are ubiquitous in slow-growing mycobacteria

In a previous bioinformatics approach aimed at finding genes that were horizontally acquired during the evolution of Mtb, Veyrier et al described *rv1722* as a gene that was likely acquired at the point where fast-growing mycobacteria evolved into slow-growing mycobacteria(49). To confirm this finding, we first performed an extensive protein BLAST analysis, which suggested that Rv1722 homologs are indeed predominantly present in slow-growing mycobacteria (Table S1).

To visualize the distribution of *rv1722* homologs throughout the Mycobacterial genus we then generated a phylogenetic tree containing Mtb and 227 non-tuberculous mycobacteria (NTM) type strains, including fast-growing and slow-growing mycobacteria, onto which we mapped the presence of *rv1722* homologs (Fig 7). Homologs of *rv1722* were present in 85 NTM type strains, with the lowest sequence identity above our 50% cutoff found in *M. parakoreensis* (78%).

**Figure 7.**
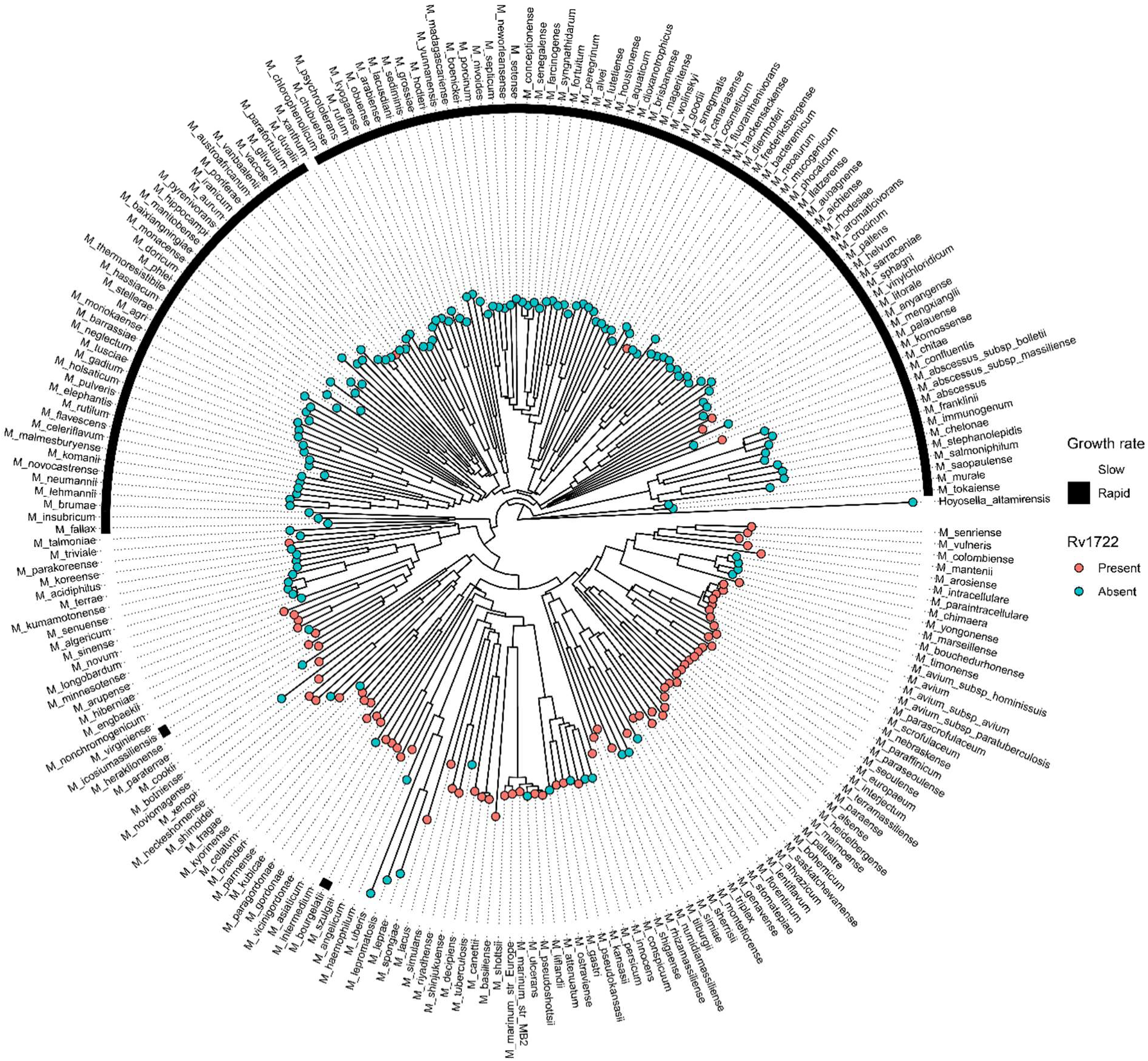
Rv1722 homologs are predominantly present in slow-growing mycobacteria. Phylogenetic tree showing the presence of rv1722 homologs and growth speed across mycobacteria. The phylogenetic tree was constructed with the full genome sequences of Mtb, 227 NTM type strains, and contains *H. altamirensis* as outgroup. The presence of rv1722 homologs was assessed using NCBI Nucleotide BLAST, with a 50% sequence similarity cutoff.

This tree corroborates that *rv1722* homologs are predominantly present in slow-growing mycobacteria, but also revealed noticeable exceptions of fast-growing mycobacteria containing an *rv1722* homolog and slow-growing mycobacteria lacking an *rv1722* homolog. Importantly, our genetic analysis results are in line with the absence of GABA-trehalose in the fast-growing mycobacteria *M. abscessus* and *M. smegmatis* exposed to stress conditions (Fig 1A). Taken together, our phylogenetic analyses show that *rv1722* is enriched in the genomes of slow-growing mycobacteria.

## Discussion

Previous work has shown that metabolic adaptation is key for the virulence of Mtb, but the current knowledge on these adaptations is rooted in gene-centric approaches that first identify metabolic enzymes of interest based on sequence homology and then confirm their function. While powerful, homology-based methods are intrinsically biased towards conserved enzymes of known function and fail to identify the most novel, organism-specific metabolism. We applied a bottom-up approach in which we first screened for unknown metabolites involved in metabolic adaptation and then identified their enzymatic origin. In doing so, we revealed GABA-trehalose, a previously unrecognized metabolite that is rapidly produced at millimolar levels by Mtb exposed to phagosome-like stresses. In addition, we show that GABA-trehalose is synthesized from GABA and trehalose via an ATP-dependent reaction that is catalyzed by the uncharacterized enzyme Rv1722 that we name GABA-trehalose synthetase (GabtS). Our results underline that a bottom-up approach enables the discovery of metabolites and pathways that are refractory to classical approaches. Combined, these results reveal a previously unrecognized, but quantitatively dominant, pathway involved in metabolic adaptation of Mtb.

While GABA-trehalose has not been described before, its building blocks GABA and trehalose are ubiquitous and widely studied. Trehalose is present in plants, yeast and prokaryotes, where it can serve as osmolyte and carbon-storage metabolite, and plays a pivotal role in the response to stresses like drought(50, 51). In Mtb, trehalose is a part of the mycolyl glycolipid cell wall component unique to mycobacteria(52). Recently, trehalose derived from Mtb cell wall has been implicated in the response to hypoxia and the metabolism of persister-like bacilli, during which it serves as carbon source for anticipated cell growth(2, 53). Like trehalose, GABA is also present in many eukaryotes and prokaryotes. While it serves as signal metabolite and increases stress tolerance in plants, and as a neurotransmitter in humans, its function in microorganisms is less clear(54). Nevertheless, GABA has been associated with microbial acid tolerance and signaling(55, 56). GABA is an intermediate in the GABA-shunt, but the function of the GABA-shunt in Mtb is similarly unclear. Still, it has been hypothesized to serve as a shunt of the TCA cycle, an entry mechanism for alternative carbon sources into the TCA cycle(57), or as a proton sink for acid stress protection(58). Recent studies have shown that it is actively involved in Mtb central carbon metabolism and able to drive succinate production(59, 60). Here, we show that GABA plays a prominent role under stress conditions and link it to trehalose metabolism via a novel pathway.

Given that GABA-trehalose is produced at millimolar levels in stressed Mtb, the pathway leading to GABA-trehalose formation likely plays an in important role in the metabolic adaptation to stress. Previous studies on Mtb have shown that adaptation to hypoxia leads to succinate accumulation and excretion(7, 46). This adaptation enables Mtb to run a partial reductive TCA cycle and maintain an energized membrane. Our results show that hypoxia, and also nitrosative stress and a combination thereof, not only lead to succinate accumulation, but also to GABA accumulation and excretion. Both hypoxia and NO inhibit oxidative phosphorylation, leading to a severe increase in the NADH/NAD+ ratio, decrease in the ATP/ADP ratio, and TCA cycle remodeling(46). We hypothesize that the increased NADH/NAD+ ratio pushes the first NADH-dependent step of the GABA-shunt forward, while the accumulation of succinate and increased NADH/NAD+ ratio together inhibit the final NAD+-dependent step of the GABA-shunt, ultimately leading to GABA accumulation. Our data show that accumulated GABA is not only excreted like succinate, but also converted into GABA-trehalose by Rv1722. We postulate that these mechanisms may mitigate potentially toxic GABA accumulation while providing thermodynamically favorable conditions for the continuous production of GABA, which regenerates NAD+ via the first step of the shunt and mitigates the increased NADH/NAD+ ratio.

Despite its role in the metabolic response to hypoxia and acid-nitrosative stress, the gene *rv1722* is not part of the stress-responsive dosR regulon and not commonly upregulated under stress conditions(61). Moreover, the protein Rv1722 has been detected in non-stressed Mtb(62). This constitutive expression suggest that GABA-trehalose formation is not regulated at the gene- or protein-level. In line with this, we show that GABA-trehalose formation is driven by GABA accumulation. Constitutive expression of Rv1772 enables a very rapid response to suddenly increasing GABA levels, which could be beneficial for rapid stress adaptation. Although GABA excretion and GABA-trehalose synthesis both mitigate GABA accumulation, GABA-trehalose does not leave the cell and potentially functions as a metabolic reservoir for carbon and nitrogen to be used when the redox balance is restored. Gillet et al recently suggested that *Mycobacterium smegmatis* similarly stocks carbon in the form of glycerides during adaptation to hypoxia(63). However, stocking GABA-trehalose comes at a price as it requires ATP, while ATP levels drop over 3-fold under hypoxia(45, 46).

Metabolism aside, we annotated Rv1722 as GABA-trehalose synthetase. Rv1722 belongs to the superfamily of ATP-grasp enzymes and has been bioinformatically annotated as potential (biotin) carboxylase and as glutathione synthetase ATP-binding domain-like protein(64–66). Our biochemical results now confirm that it functions as a ligase. However, unlike other ATP-grasp enzymes, it does not catalyze the ligation of an amine or thiol and a carboxylic acid, but the ligation of a hydroxyl and a carboxylic acid(67). Our results thus reveal Rv1722 as a non-canonical ATP-grasp enzyme.

Our biochemical (Fig 2) and phylogenetic results (Fig 7) show that *rv1722* and its homologs are predominantly present in slow-growing mycobacteria. Slow-growing mycobacteria have evolved from the older fast-growing mycobacteria in a major genetic shift, which reportedly introduced *rv1722* through horizontal gene transfer(49). Several publications argue that horizontal gene transfers shaped the virulence of the slow-growing mycobacteria that ultimately evolved into the MTB complex(68–71). Nevertheless, gene essentiality screens predict *rv1722* to be non-essential(72). Similar to our work, Layre et al discovered that the horizontally acquired genes *rv3377c* and *rv3378c* are responsible for the production of the Mtb-specific metabolite 1-tuberculosinyladenosine (TbAd)(15). While *rv3377c-rv3378c* are also predicted to be non-essential by most genetic screens, TbAd was later shown to prevent phagolysome acidification(73), a key process in the antimicrobial activity of macrophages. The function of GABA-trehalose formation remains to be determined.

Concluding, we used a bottom-up metabolite-to-gene approach to discover a new stress response pathway in Mtb. This pathway is a unique extension of Mtb central carbon metabolism and provides a fundamental new insight in Mtb’s response to immune-stresses, while GABA-trehalose holds promise as a pathogen-specific biomarker. Several questions remain to be answered, however. Future research could, for example, focus on the breakdown pathway of GABA-trehalose and the phenotype of a *rv1722* knockout. Since GABA-trehalose appears to be unique to mycobacteria, its biosynthesis could potentially serve as a selective target to treat TB, a disease that still kills more than a million people per year.

## Acknowledgements

The authors would like to thank Kyu Rhee and Piet Borst for a critical reading of the manuscript and Huub Op den Camp, Inna Krieger and Valerie Koeken for discussions. We are thankful to Ben Marais and Nathan Bachman for sharing phylogenetic data of mycobacterial species.

