## Supplemental Information for "A new stress-response pathway in *Mycobacterium tuberculosis*"

### Supplementary Information

#### Methods

##### Macrophage infection

Whole blood from three healthy volunteers was obtained from Sanquin Bloodbank (Nijmegen, the Netherlands), as approved by the Ethics Committee of Radboud University Medical Center (Nijmegen, the Netherlands). Peripheral blood mononuclear cells (PBMCs) were isolated by Ficoll-Paque density gradient centrifugation, followed by 3 wash steps with cold phosphate-buffered saline (PBS) and resuspension in Dutch-modified RPMI 1640 medium (Invitrogen) supplemented with 50 µg/mL gentamicin, 2 mM Glutamax (GIBCO), and 1 mM pyruvate (GIBCO). PBMCs of each donor were seeded in 10 cm Petri dishes (Corning) at $40x10^6$ cells/dish and incubated at 37°C in 5% $CO_2$ in a humidified incubator (the same conditions were used throughout this experiment) to allow monocyte purification through plastic adhesion. After 2 hours, non-adherent lymphocytes were removed by carefully washing the plates three times with pre-warmed PBS (37°C), followed by the addition of 10 mL pre-warmed differentiation medium (complete RPMI supplemented with 10% (v/v) human pooled serum and 5 ng/mL macrophage colony-stimulating factor (M-CSF, Miltenyi Biotec) and incubation. On day 3 after seeding, 2/3$^{rd}$ of the spent differentiation medium was replaced with fresh pre-warmed differentiation medium. On day 7, the cells were washed three times with pre-warmed PBS and harvested by adding 4 mL Versene (Gibco), incubating for 30 min at 37°C, and collecting the cell suspension. Remaining adherent cells were scraped twice and collected in the same tube. The cell suspensions were centrifuged at 1700 rpm for 10 min at room temperature and resuspended in complete RPMI to a final density of $1x10^6$ cells/mL. To activate the cells, 1 mL of the cell suspensions were diluted into 1 mL antibiotic-free medium with or without supplementation of 200 ng human IFN-γ 1-B (IMMUKINE, Clinigen Healthcare B.V., Schiphol, The Netherlands) in 6-well plates, followed by incubation overnight. On day 8, the human monocyte-derived macrophages (hMDMs) were infected with *M. tuberculosis*. To achieve this, first, *M. tuberculosis* H37Rv bacteria were harvested by centrifuging a liquid culture for 5 min at room temperature and 5,000 g and resuspended in 1 mL 0.05% (v/v) Tween 20 in PBS. This step was repeated once, followed by passaging the cell suspension 10 times through a 27-gauge needle to dislodge aggregates. The *M. tuberculosis* H37Rv bacteria were then diluted to a final concentration of $0.5x10^6$ CFU/mL for a multiplicity of infection of 1 (MOI=1), $2.5x10^6$ CFU/mL (MOI=5), and $5.0x10^6$ CFU/mL (MOI=10) in complete RPMI with 10% (v/v) HPS and 100 ng/mL IFN-γ 1-B. Then, medium was removed from the hMDM-containing wells and 2 mL of the *M. tuberculosis* H37Rv suspension was added, followed by incubation. After 4 hours, medium was removed, and cells were washed three times with pre-warmed PBS to remove extracellular bacteria. To extract the metabolites, 1 mL of quenching buffer (20% (v/v) ultrapure water (Veolia water solutions & technologies, High Wycombe, UK), 40% (v/v) acetonitrile and 40% (v/v) methanol) was added to each well, except for the hMDM-free control, followed by incubation on an ice-pack for 10 min. Next, wells were scraped, and the suspension was transferred to 2 mL screw cap tubes containing 0.1 mm zirconia beads, and bead-beaten as described above. The bacteria in the medium of the hMDM-free control were centrifuged for 10 min at room temperature and 4000 g, resuspended in quenching buffer, and then subjected to bead-beating as the hMDM-containing samples. Finally, the metabolite extracts were filter-sterilized with 0.22 µm Spin-X centrifuge tube filters (Costar, Corning, New York, USA) and stored at -20°C until metabolomics analysis.

#### Synthesis of GABA-trehalose, GABA-glucose and $^{13}$C2-GABA

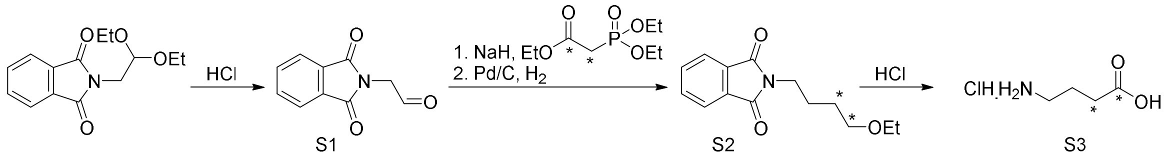


N-phthalimidyl-2-aminoacetaldehyde (S1)

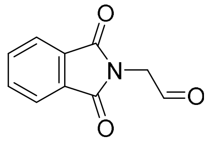

2-(2,2-diethoxyethyl)isoindoline-1,3-dione (2.0 g, 7.6 mmol) was dissolved in 1 M HCl, heated to reflux and stirred for 1 h. After complete conversion, the solution was cooled to room temperature, extracted with DCM. The resulting organic layer was washed with sat. aq. NaCl, dried over $MgSO_4$ and concentrated *in vacuo*. The product was obtained as a solid in 89% yield (1.3 g, 6.9 mmol). $R_F$ 0.20 (1:1 EtOAc:heptane); $^1$H NMR (500 MHz, Chloroform-*d*) δ 9.65 (s, 1H), 7.89 (dd, *J* = 5.5, 3.1 Hz, 2H), 7.76 (dd, *J* = 5.5, 3.0 Hz, 2H), 4.56 (s, 2H); $^{13}$C NMR (126 MHz, $CDCl_3$) δ 193.7, 167.7, 134.5, 132.0, 123.8, 47.5.

$^{13}C_2$-ethyl 4-phthalimidobutyrate (S2)

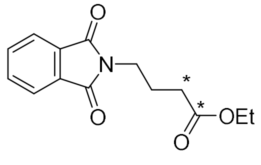

A solution of $^{13}C_2$-triethyl phosphonoacetate (800 µL, 4.0 mmol) in THF (30 mL) was cooled to -78 °C. To this solution, solid NaH (196 mg, 4.9 mmol, 60 wt%) was added. The mixture was allowed to warm to 0 °C and kept at that temperature for 20 min. The solution was cooled to -78 °C and a solution of S1 (1.27 g, 6.7 mmol) in THF (30 mL) was dropwise added. The solution was allowed to slowly warm to -20 °C. After 30 mins, complete conversion was observed ($R_F$ 0.15 (1:4 EtOAc:heptane)). An sat. aq. solution of $NaHCO_3$ (60 mL) was added, the solution was diluted with EtOAc and the layers were separated. The organic layer was dried over $MgSO_4$ and concentrated *in vacuo*. The product was purified by column chromatography (1:4 EtOAc:heptane) to yield a white solid (940 mg, 3.6 mmol, 90%). The product was dissolved in EtOH (10 mL), purged with argon. A catalytic amount of Pd/C (10 wt% Pd) was added. The atmosphere was exchange to that of $H_2$. After complete conversion, the solution was purged with argon, the suspension filtered over celite and the filtrate concentrated *in vacuo*. The product was obtained as white solid in 90% yield over two steps (943 mg, 3.6 mmol). $R_F$ 0.15 (1:4 EtOAc:heptane); $^1$H NMR (500 MHz, CHLOROFORM-*D*) δ 7.90 – 7.81 (m, 2H), 7.74 – 7.68 (m, 2H), 4.10 (qd, *J* = 7.1, 3.0 Hz, 2H), 3.75 (td, *J* = 6.9, 4.3 Hz, 2H), 2.37 (dq, *J* = 128.0, 7.5 Hz, 2H), 2.03 (ddt, *J* = 9.0, 7.3, 3.6 Hz, 2H), 1.24 (td, *J* = 7.2, 1.5 Hz, 3H); $^{13}$C NMR (126 MHz, CHLOROFORM-*D*) δ 172.7 (d, *J* = 58.2 Hz), 168.4, 134.0, 132.2, 123.3, 60.6 (d, *J* = 3.9 Hz), 37.3 (d, *J* = 4.7 Hz), 31.7 (d, *J* = 58.0 Hz), 25.49 – 23.14 (m), 14.3 (d, *J* = 3.1 Hz).

$^{13}C_2$ GABA (S3)

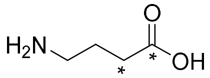

A mixture of S2 (940 mg, 3.6 mmol) was dissolved in 3 M HCl (20 mL) and heated to a reflux for 12 h. The clear solution was cooled with an ice-bath and precipitation was observed. The precipitate, phthalic acid, was filtered off. The filtrate was concentrated *in vacuo*. The solid was resuspended in water, filtered and concentrated *in vacuo.* The resuspension in water and filtration procedure was repeated thrice more. The product was obtained as a white solid in 68% yield (360 mg, 2.6 mmol), contaminated with 12 wt% phthalic acid. $^{1}H$ NMR (500 MHz, $D_2O$) δ 2.94 (dq, *J* = 8.7, 3.3 Hz, 2H), 2.41 (dq, *J* = 128.7, 7.2 Hz, 2H), 1.85 (dtd, *J* = 12.2, 7.3, 2.5 Hz, 2H); $^{13}C$ NMR (126 MHz, $D_2O$) δ 177.1 (d, *J* = 54.5 Hz), 38.8 (d, *J* = 4.8 Hz), 30.7 (d, *J* = 54.2 Hz), 22.1 (d, *J* = 32.6 Hz).

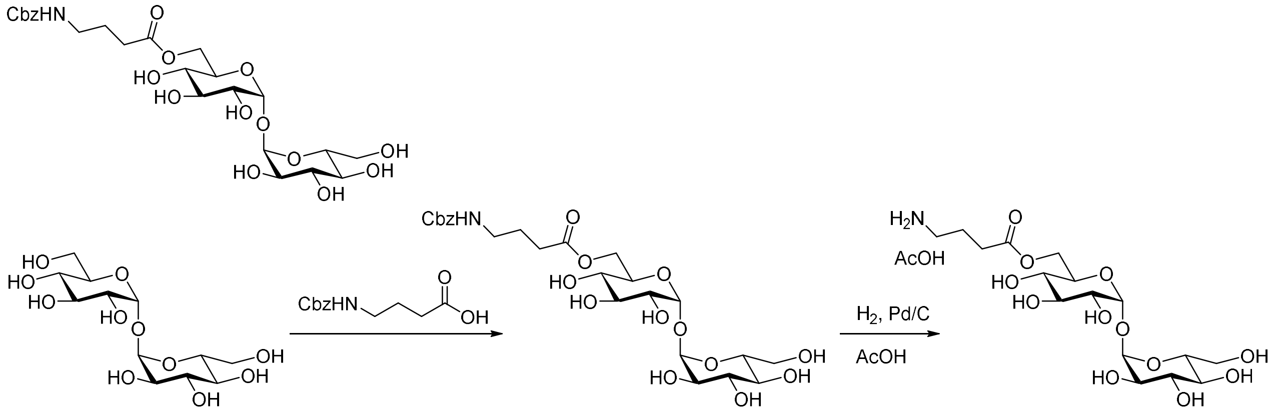


Threalose S4 S5

6'-*N*CbzGABA-Trehalose (S4)

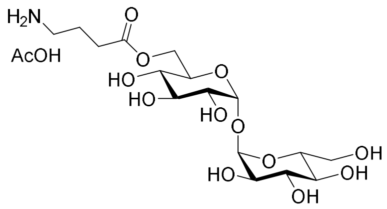


A solution of Trehalose-dihydrate (690 mg, 1.8 mmol) was co-evaporated twice with dry pyridine, and redissolved in dry DMF (10 mL). This solution was added to a solution of *N*-CbzGABA (476 mg, 2.0 mmol) and HATU (763 mg, 2.0 mmol) which has been stirred for 30 minutes at rt in dry pyridine (10 mL). The combined solution was left standing for 36 h. The solution was concentrated *in vacuo*. The product was purified twice by column chromatography (1:4 $H_2O$:MeCN) to yield a white solid in 15% yield (157 mg, 1.82 mmol). $R_F$ 0.58 (1:4 $H_2O$:MeCN); $^{1}H$ NMR (500 MHz, DMSO) δ 7.42 – 7.22 (m, 6H), 5.05 (d, *J* = 5.4 Hz, 1H), 5.01 (s, 2H), 4.88 – 4.85 (m, 2H), 4.84 (d, *J* = 3.6 Hz, 1H), 4.76 (dd, *J* = 5.1, 2.0 Hz, 2H), 4.68 (t, *J* = 6.0 Hz, 2H), 4.35 (t, *J* = 6.0 Hz, 1H), 4.23 (dd, *J* = 11.7, 2.1 Hz, 1H), 4.04 (dd, *J* = 11.7, 5.6 Hz, 1H), 3.90 (ddd, *J* = 10.1, 5.5, 2.1 Hz, 1H), 3.65 (ddd, *J* = 9.9, 4.9, 2.3 Hz, 1H), 3.58 – 3.51 (m, 3H), 3.50 – 3.44 (m, 1H), 3.29 – 3.22 (m, 2H), 3.16 – 3.09 (m, 2H), 3.02 (q, *J* = 6.6 Hz, 2H), 2.30 (dd, *J* = 8.1, 7.0 Hz, 2H), 1.65 (p, *J* = 7.3 Hz, 2H); $^{13}C$ NMR (126 MHz, DMSO) δ 172.5, 156.2, 137.2, 128.4, 127.8, 127.8, 93.4, 93.3, 72.81, 72.78, 72.6, 71.53, 71.45, 70.1, 69.6, 65.2, 63.3, 60.7, 30.9, 24.8; HRMS (m/z): $[M+H]^+$ calcd for $C_{24}H_{35}NO_{14}H$: 562.2130; found: 562.2124.

6'-GABA-Trehalose

A solution of S4 (39.3 mg, 70 µmol) in dioxane (2 mL) and AcOH (10 µL, 120 µmol) was purged with argon. A catalytic amount of Pd/C (10 wt% Pd) was added. The atmosphere was exchange to that of $H_2$. After complete conversion, the solution was purged with argon, the suspension filtered over celite and lyophilized twice to remove all traces of dioxane, to yield a fluffy white solid in 95% yield, contaminated with trehalose (34.1 mg, 67 µmol). $^{1}H$ NMR (500 MHz, $D_2O$) δ 5.06 (d, *J* = 3.8 Hz, 1H), 5.03 (d, *J* = 3.8 Hz, 1H), 4.33 (dd, *J* = 12.2, 2.3 Hz, 1H),

4.21 (dd, *J* = 12.2, 5.1 Hz, 1H), 3.91 (ddd, *J* = 10.2, 4.9, 2.3 Hz, 1H), 3.80 – 3.61 (m, 5H), 3.57 – 3.48 (m, 2H), 3.40 – 3.29 (m, 2H), 2.97 – 2.88 (m, 2H), 2.46 (t, *J* = 7.3 Hz, 2H), 1.92 – 1.83 (m, 2H); $^{13}C$ NMR (126 MHz, $D_2O$) δ 174.60, 93.63, 93.45, 72.61, 72.30, 71.06, 70.98, 69.95, 69.72, 63.44, 60.52, 38.72, 30.56, 22.04; HRMS (m/z): $[M+H]^+$ calcd for $C_{16}H_{29}NO_{12}H$: 428.1763; found: 428.1779.

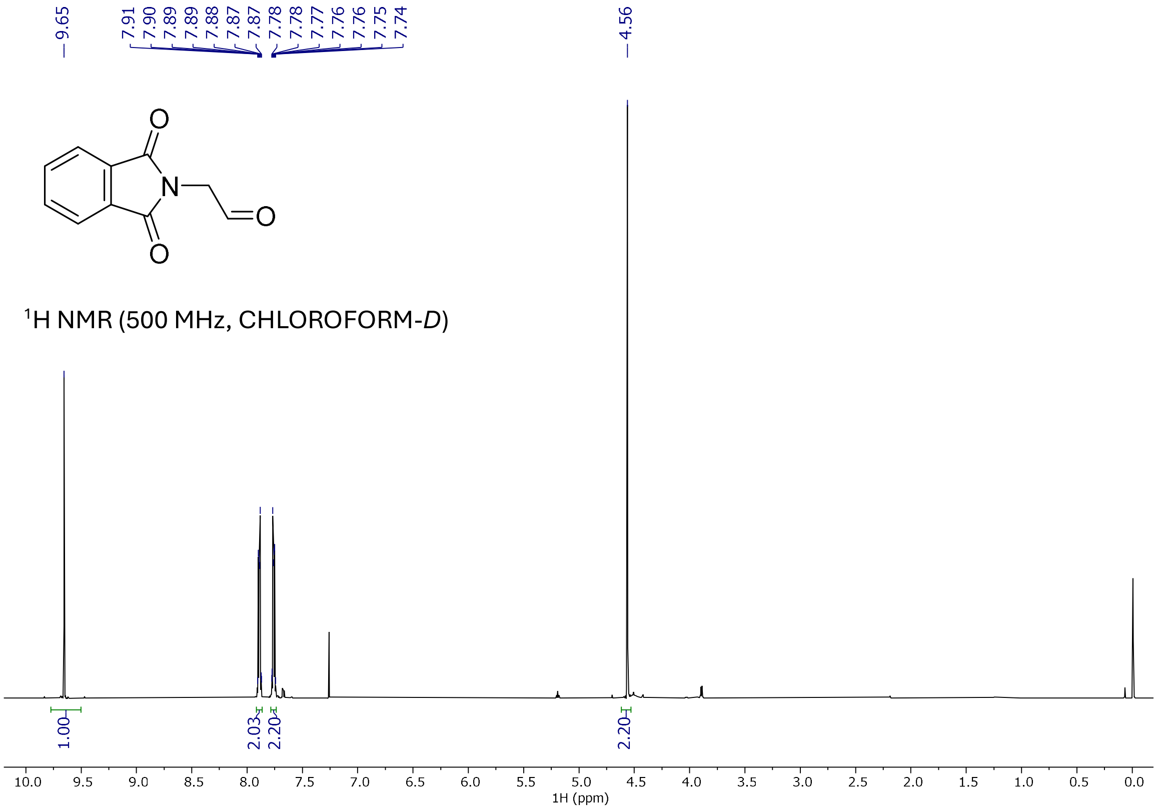

9.65
7.91 7.90 7.89 7.89 7.88 7.87 7.87 7.78 7.78 7.77 7.76 7.76 7.75 7.74
4.56
O
N
O
O
1H NMR (500 MHz, CHLOROFORM-D)
1.00
2.03
2.20
2.20
10.0 9.5 9.0 8.5 8.0 7.5 7.0 6.5 6.0 5.5 5.0 4.5 4.0 3.5 3.0 2.5 2.0 1.5 1.0 0.5 0.0
1H (ppm)


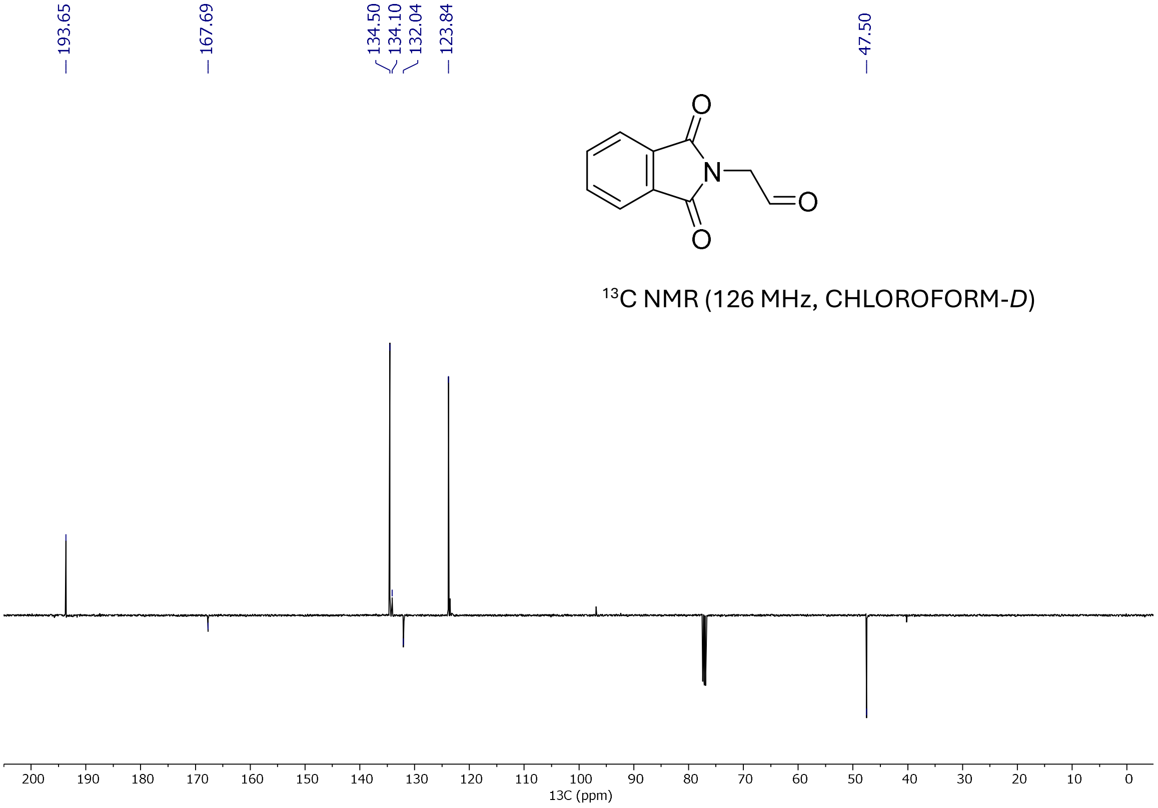

193.65
167.69
134.50 134.10 132.04
123.84
47.50
O
N
O
O
13C NMR (126 MHz, CHLOROFORM-D)
200 190 180 170 160 150 140 130 120 110 100 90 80 70 60 50 40 30 20 10 0
13C (ppm)

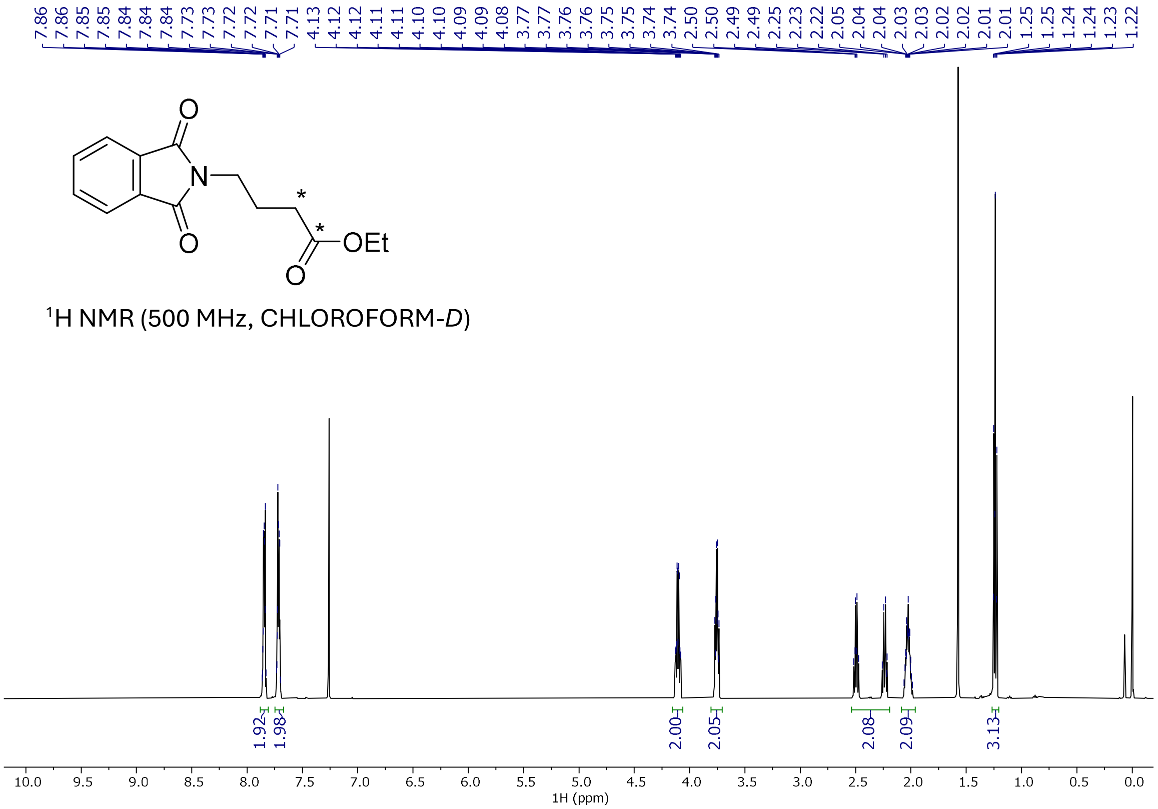

7.86 7.86 7.85 7.85 7.84 7.84 7.84 7.73 7.73 7.72 7.72 7.71 7.71 4.13 4.12 4.12 4.11 4.11 4.10 4.10 4.09 4.09 4.08 3.77 3.77 3.76 3.76 3.75 3.75 3.74 3.74 2.50 2.50 2.49 2.25 2.23 2.22 2.05 2.04 2.04 2.03 2.03 2.02 2.02 2.01 2.01 1.25 1.25 1.24 1.24 1.23 1.22
O
N
O
*
*
OEt
O
$^{1}$H NMR (500 MHz, CHLOROFORM-*D*)
1.92
1.98
2.00
2.05
2.08
2.09
3.13
10.0 9.5 9.0 8.5 8.0 7.5 7.0 6.5 6.0 5.5 5.0 4.5 4.0 3.5 3.0 2.5 2.0 1.5 1.0 0.5 0.0
1H (ppm)


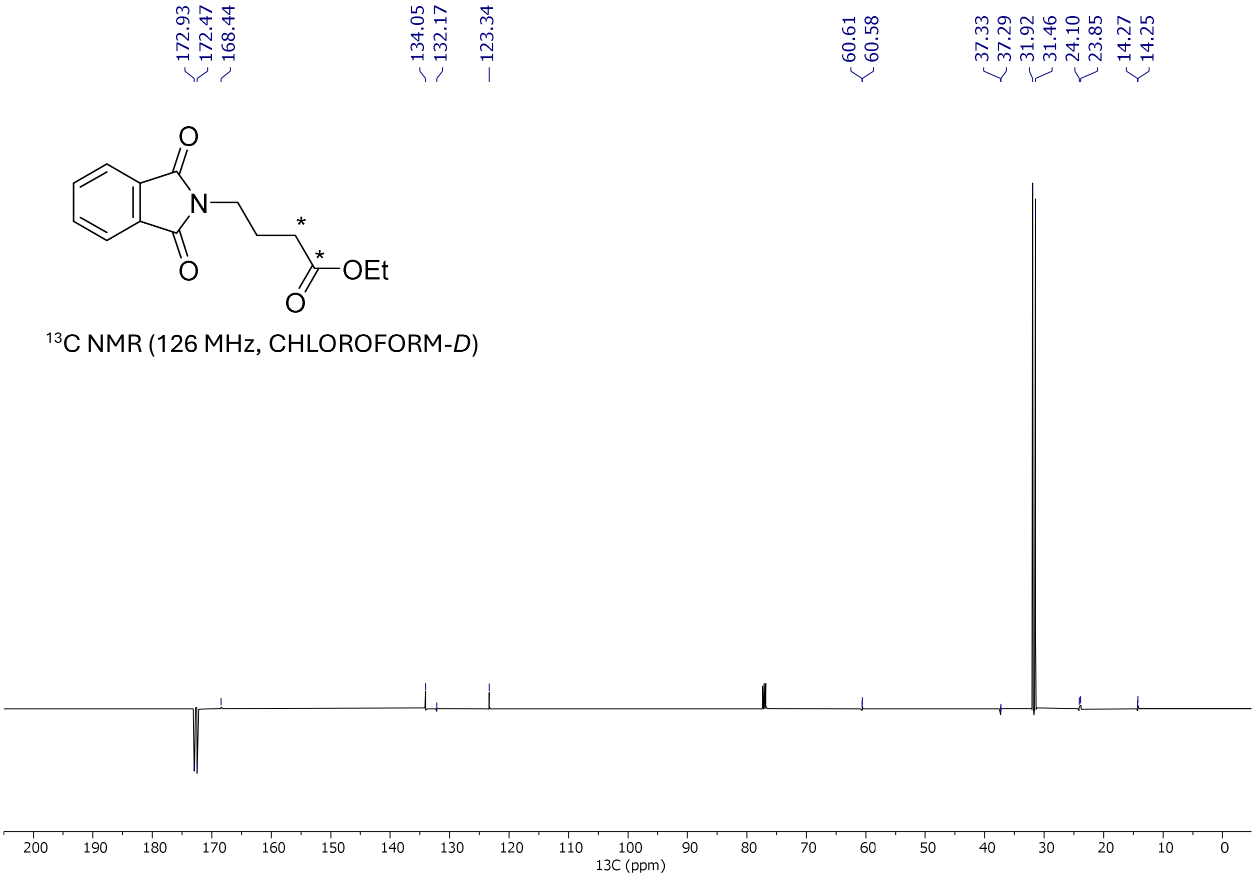

172.93 172.47 168.44
134.05 132.17
123.34
60.61 60.58
37.33 37.29 31.92 31.46 24.10 23.85 14.27 14.25
O
N
O
*
*
OEt
O
$^{13}$C NMR (126 MHz, CHLOROFORM-*D*)
200 190 180 170 160 150 140 130 120 110 100 90 80 70 60 50 40 30 20 10 0
13C (ppm)

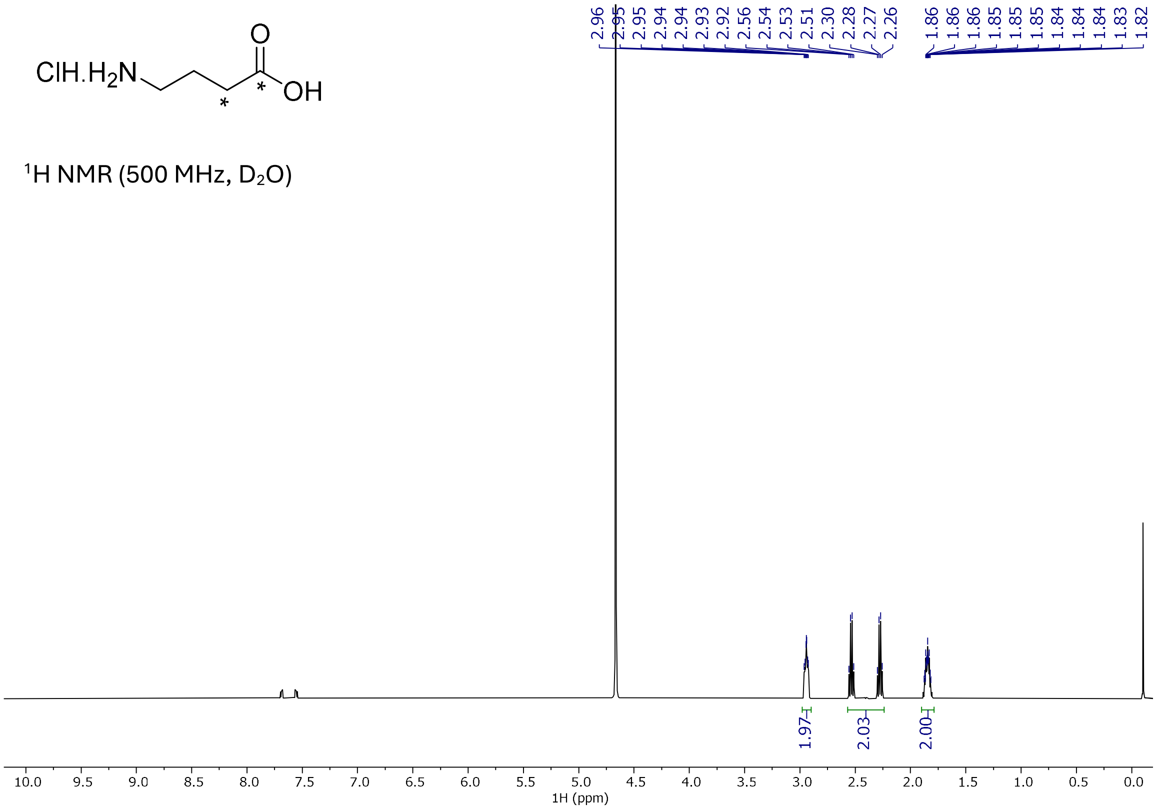


$^1$H NMR (500 MHz, $D_2O$)

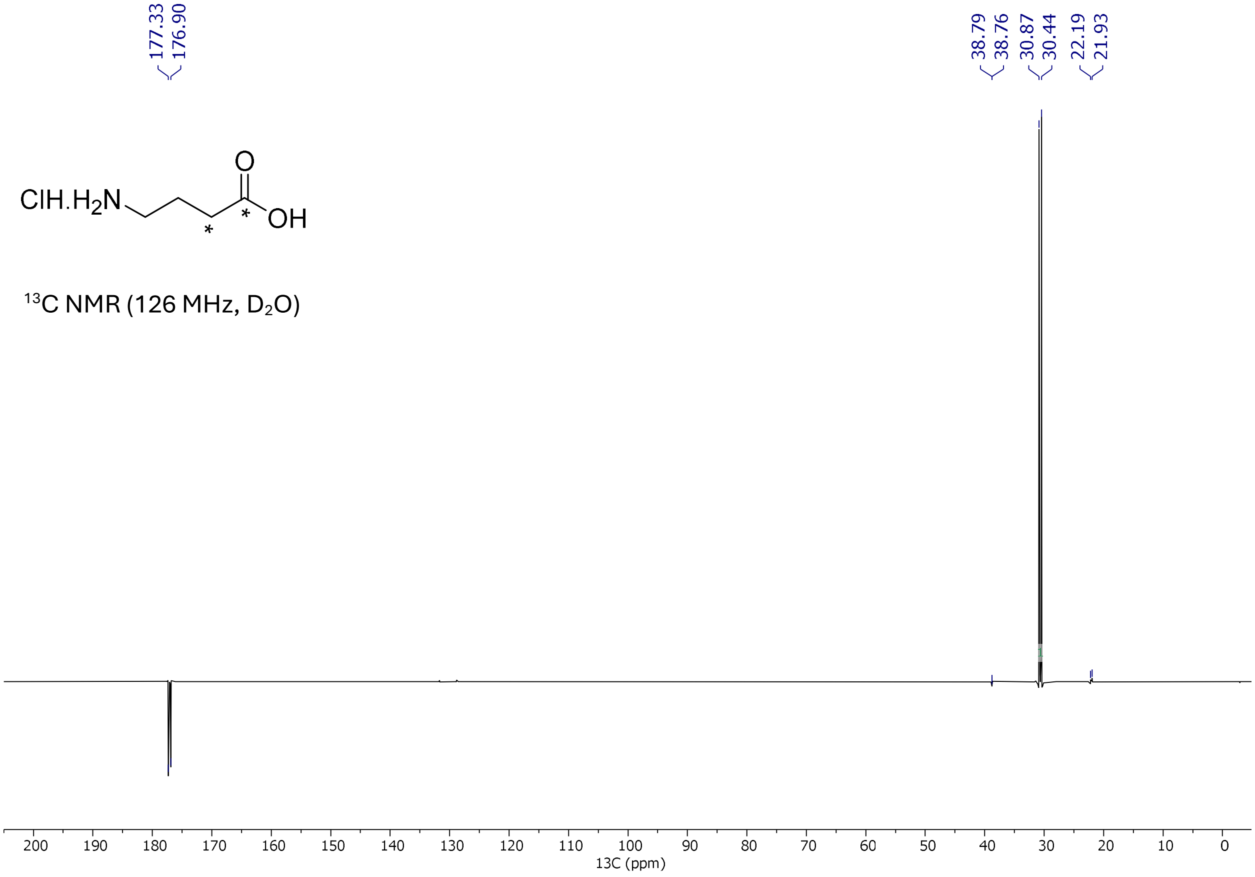


$^{13}$C NMR (126 MHz, $D_2O$)

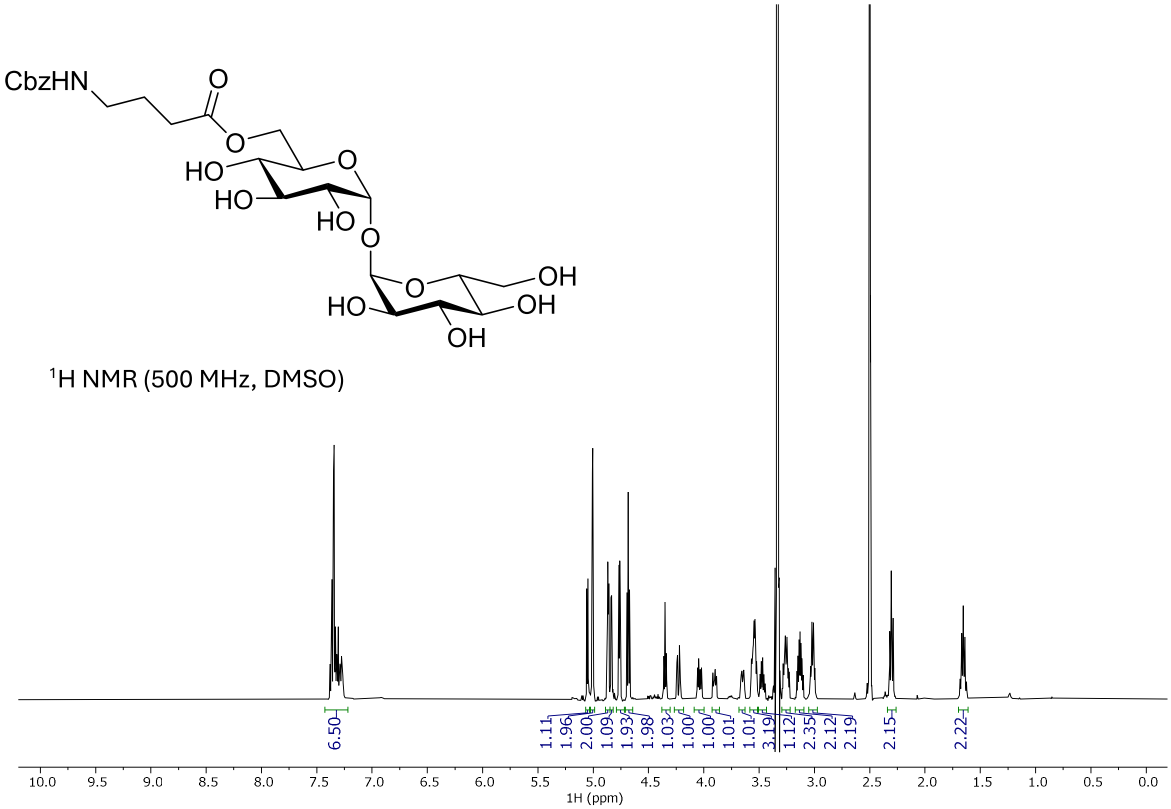
CbzHN
O
HO
HO
HO
O
O
OH
HO
OH
OH
¹H NMR (500 MHz, DMSO)
6.50
1.11
1.96
2.00
1.09
1.93
1.98
1.03
1.00
1.00
1.01
1.01
3.19
1.12
2.35
2.12
2.19
2.15
2.22
10.0 9.5 9.0 8.5 8.0 7.5 7.0 6.5 6.0 5.5 5.0 4.5 4.0 3.5 3.0 2.5 2.0 1.5 1.0 0.5 0.0
1H (ppm)

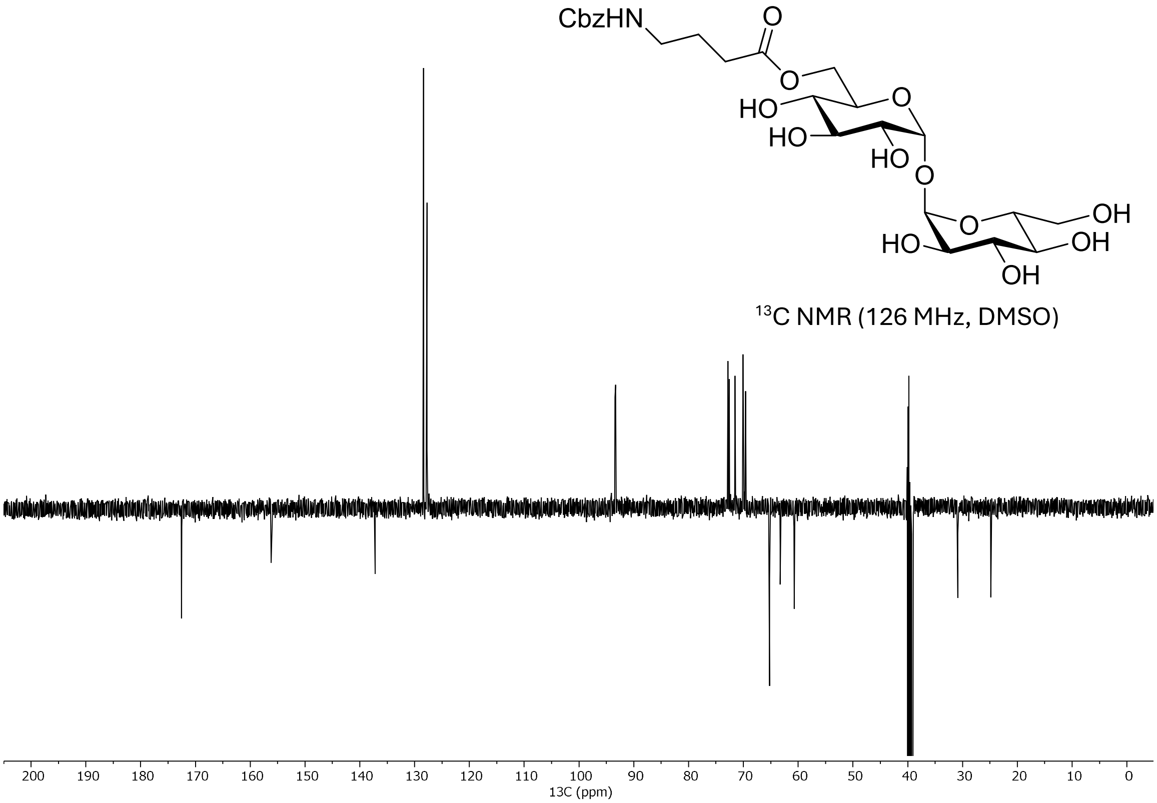
CbzHN
O
HO
HO
HO
O
O
OH
HO
OH
OH
¹³C NMR (126 MHz, DMSO)
200 190 180 170 160 150 140 130 120 110 100 90 80 70 60 50 40 30 20 10 0
13C (ppm)

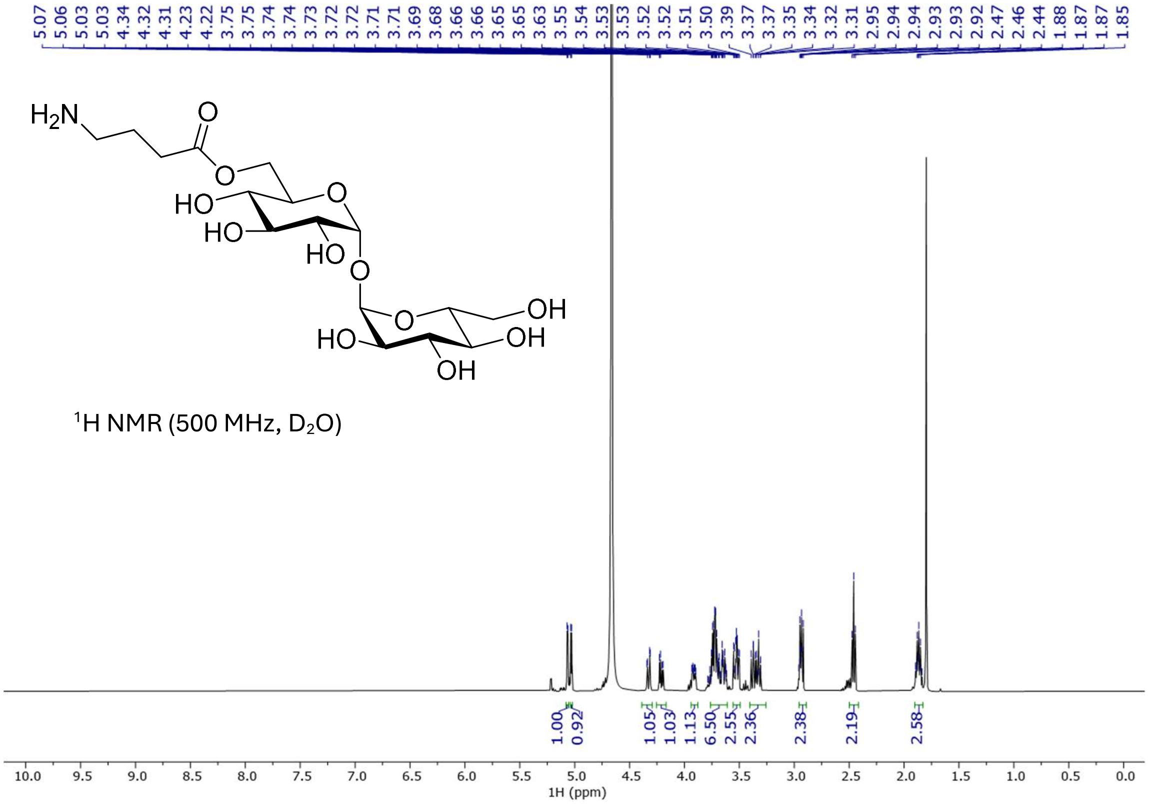
5.07 5.06 5.03 5.03 4.34 4.32 4.31 4.23 4.22 3.75 3.75 3.74 3.74 3.73 3.72 3.72 3.71 3.71 3.69 3.68 3.66 3.66 3.65 3.63 3.55 3.54 3.53 3.53 3.52 3.52 3.51 3.50 3.39 3.37 3.37 3.35 3.34 3.32 3.31 2.95 2.94 2.94 2.93 2.93 2.92 2.47 2.46 2.44 1.88 1.87 1.87 1.85
H2N
O
HO
HO
HO
OH
OH
OH
[1]H NMR (500 MHz, D2O)
1.00 0.92 1.05 1.03 1.13 6.50 2.55 2.36 2.38 2.19 2.58
10.0 9.5 9.0 8.5 8.0 7.5 7.0 6.5 6.0 5.5 5.0 4.5 4.0 3.5 3.0 2.5 2.0 1.5 1.0 0.5 0.0
1H (ppm)

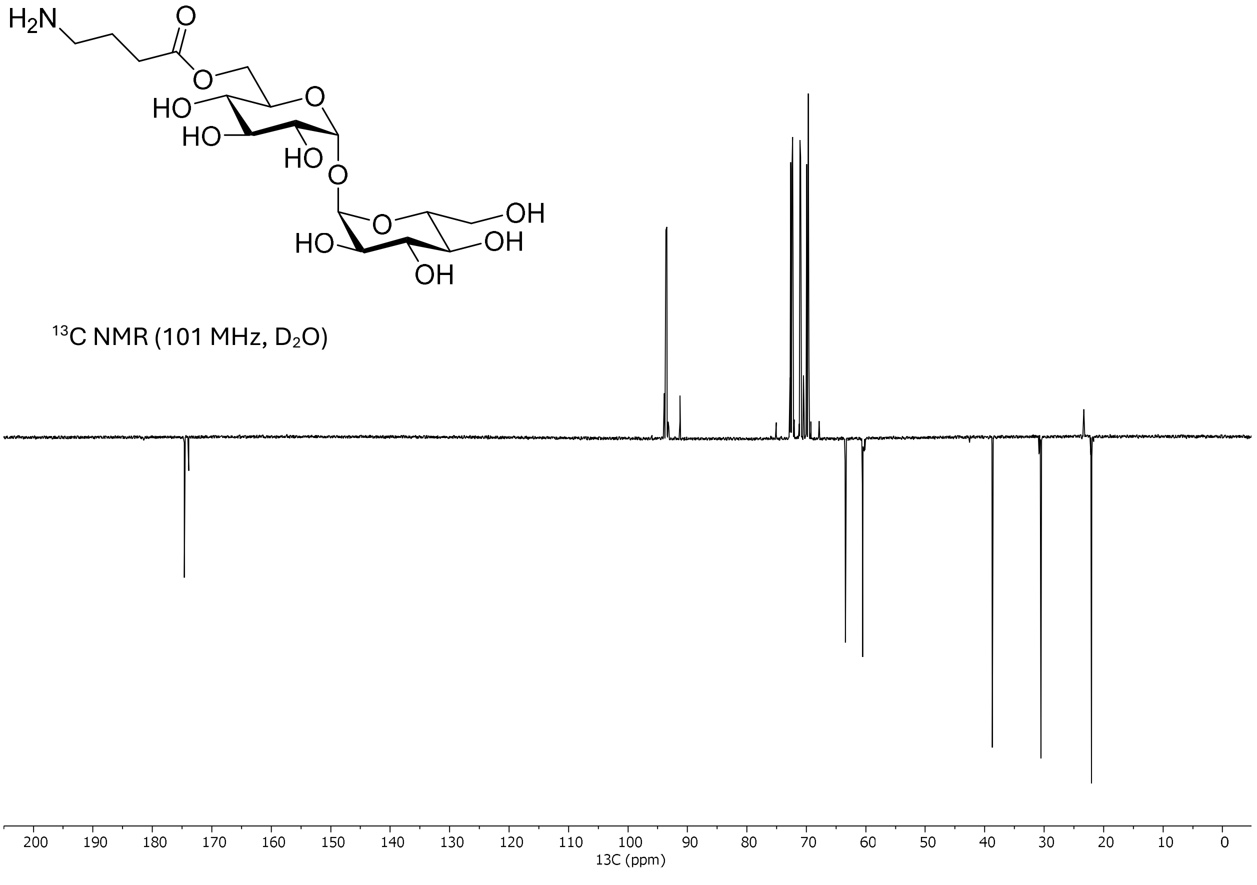
H2N
O
HO
HO
HO
OH
OH
OH
[13]C NMR (101 MHz, D2O)
200 190 180 170 160 150 140 130 120 110 100 90 80 70 60 50 40 30 20 10 0
13C (ppm)

### Figures

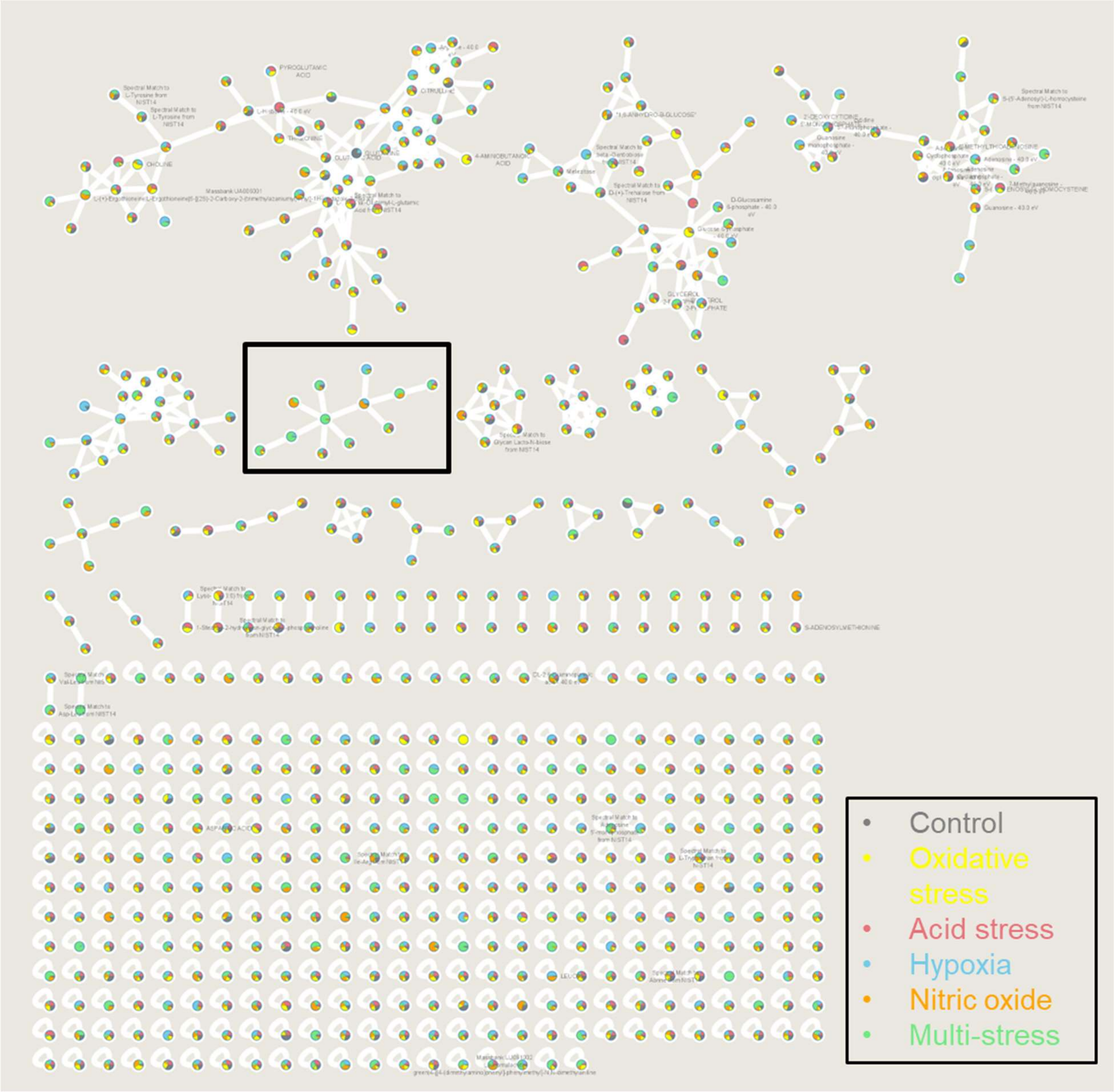


**Figure S1. Full feature-based molecular network of *Mycobacterium tuberculosis* exposed to stresses.** Each node represents a feature and is depicted as a pie chart that represents the mean relative peak area across stresses (n=3), with the same color to stress assignment as figure 1B. Nodes are connected via edges if the $MS^2$-spectra are similar (Cosine score>0.55 and more than 5 matched peaks). Nodes are putatively annotated with GNPS spectra matches. The subnetwork shown in figure 1H is highlighted by the rectangle.

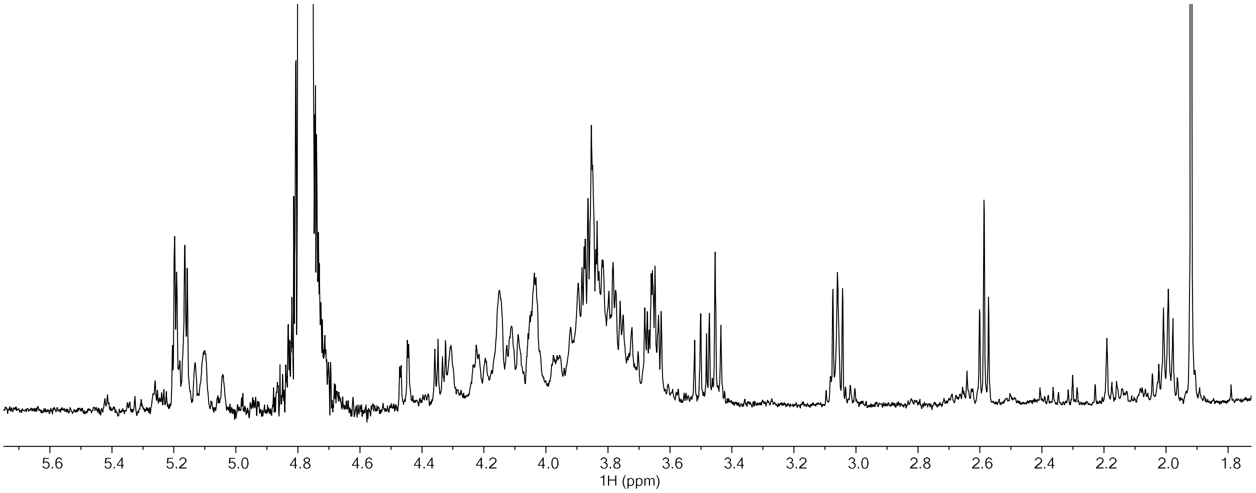


**Fig S2A. 1D 1H spectrum of concentrated extract**.

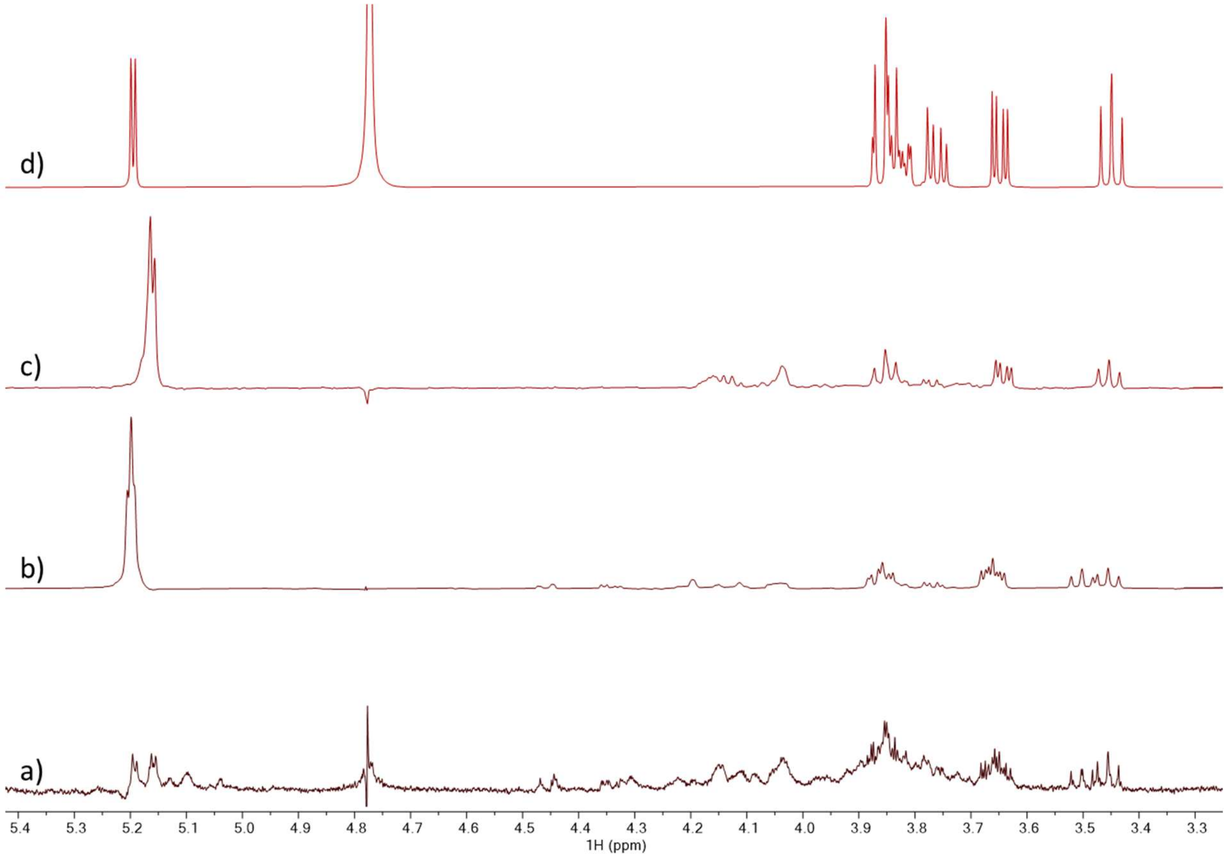


**Fig S2B**. Stacked spectra of the concentrated extract (a), selective TOCSYs of the anomeric $^{1}H$ (b,c) and trehalose (d) dissolved in $D_2O$. The H1s of the extract are slightly upfield or downfield from the trehalose H1 while the other resonances of slice “c” match trehalose well and the other slice “b” shows deviations, suggesting functionalizing of this ring.

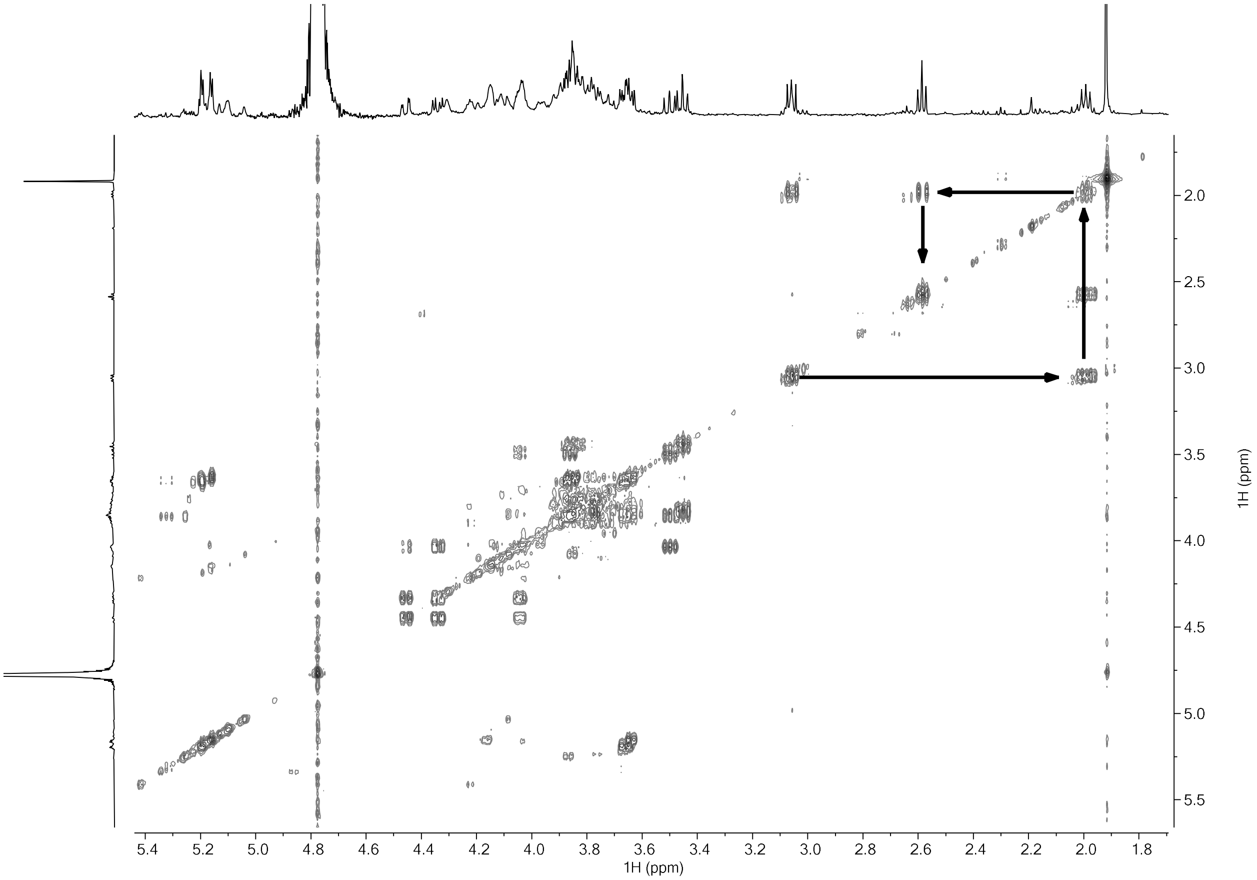


**Fig S2C. 2D COSY of concentrated extract revealing the spin network of the three $CH_2$s associated with the trehalose.** Coupling pattern and chemical shift suggested a GABA-like moiety.

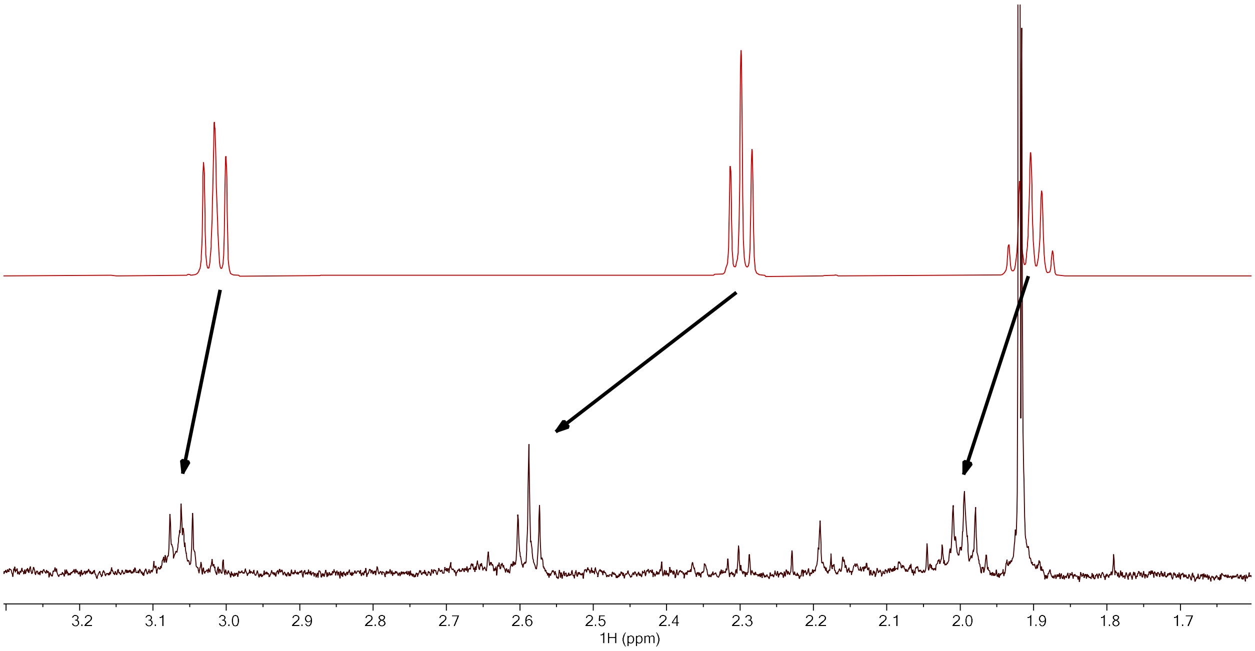

**Fig S2D. Stacked spectrum of concentrated extract (bottom) with GABA (top) indicating a modification of the pure GABA species.**

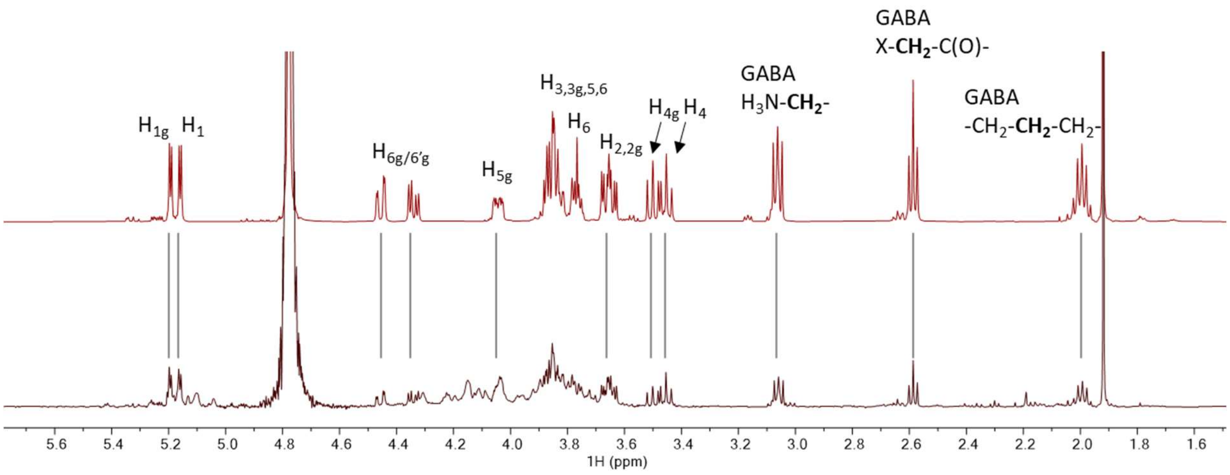


**Figure S2C. $^{1}$H 1D comparison between concentrated extract (bottom) and synthetic O6-GABA-trehalose with assignment to confirm identity.**

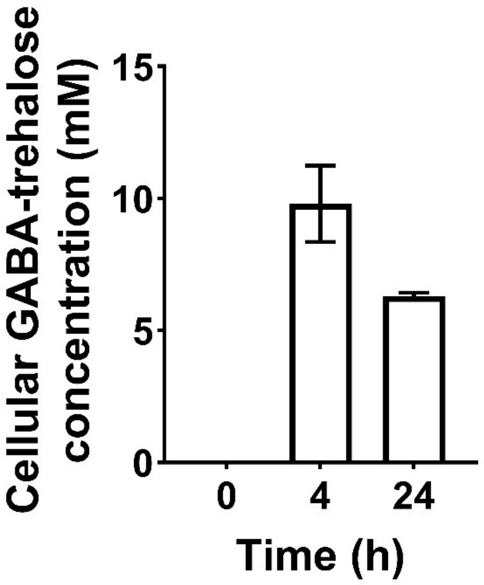


**Fig S3. Intracellular GABA-trehalose concentration in Mtb exposed to multi-stress over 24 hours.** After 4 hours of exposure to multi-stress the intracellular GABA-trehalose concentration had reached 9.8 mM and after 24 hours the concentration dropped to 6.3 mM. Data are presented as mean +/- SD, n=3.

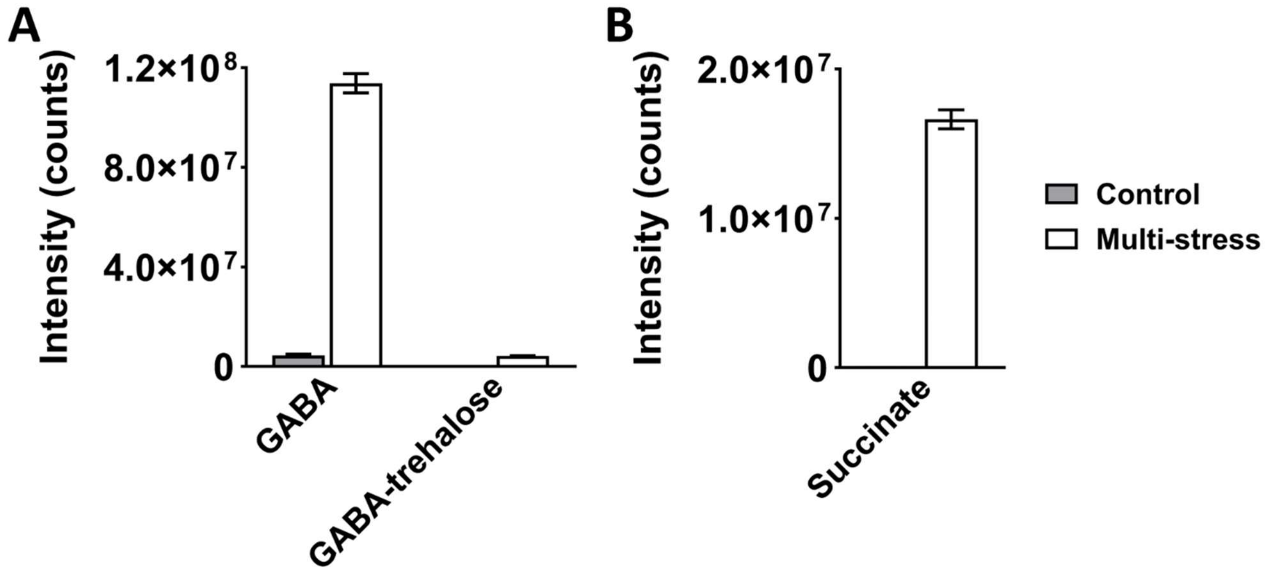


**Fig S4. GABA and succinate, but not GABA-trehalose, are excreted upon multi-stress exposure.** **A)** Intensity of GABA and GABA-trehalose in the medium of floating Mtb filters after 5 days of multi-stress exposure. **B)** Intensity of succinate in the medium of floating Mtb filters after 5 days of multi-stress exposure. Data are presented as mean +/- SD, n=3.

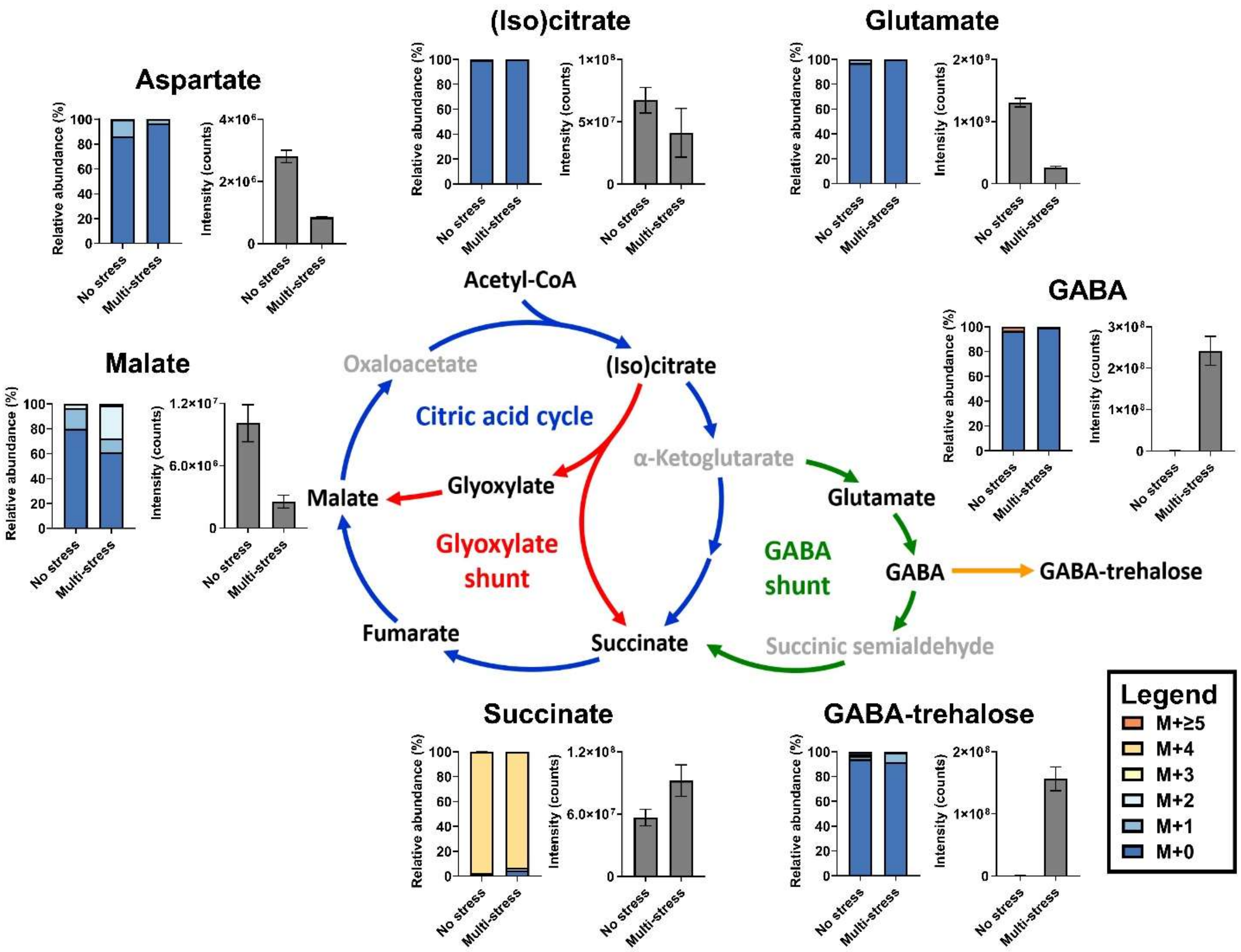


**Fig S5. The GABA-shunt is not reversed under multi-stress conditions.** Relative abundances of the isotopologs and total abundances of intermediates in central carbon metabolism 24 hours after transferring in Mtb-laden filters to medium containing $D_4$-succinate, with and without the application of multi-stress conditions. Fumarate data is not shown because succinate interfered with its isotopolog signals. Data are presented as mean (n=3) and are corrected for natural isotope abundance.

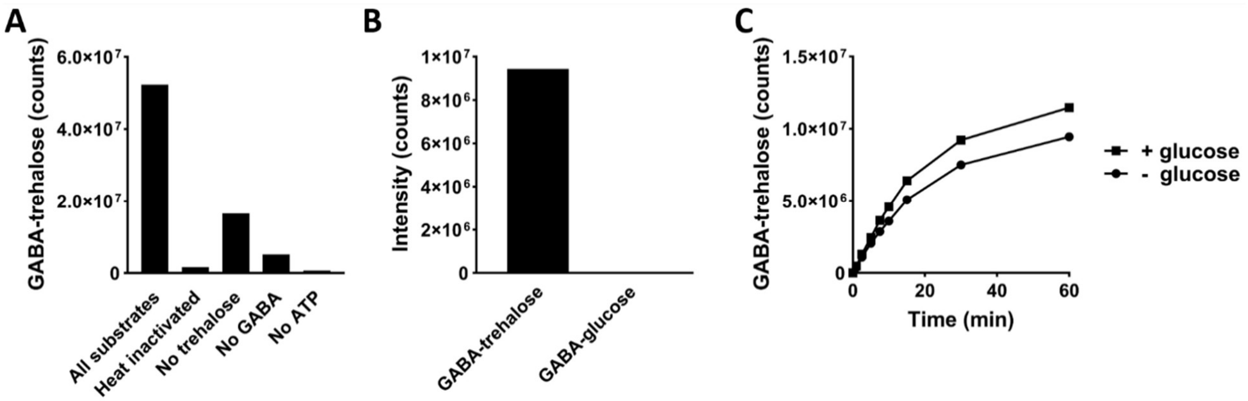


**Fig S6. Substrate screening in Mtb crude lysate, Rv1722 GABA-glucose synthesis test, and glucose competitive inhibition test. A)** Initial GABA-trehalose synthetase activity assays with Mtb crude lysate to determine the substrates required for GABA-trehalose synthesis (n=1). Remaining activity after leaving trehalose or GABA is likely caused by their presence in the crude lysate. **B)** Catalytic activity of Rv1722 towards trehalose and glucose (n=1). **C)** Enzyme assay of GABA-trehalose synthesis by Rv1722 in the presence or absence of 10 mM glucose (n=1).
